# A Telomere-to-Telomere Diploid Reference Genome and Centromere Structure of the Chinese Quartet

**DOI:** 10.1101/2025.03.30.646227

**Authors:** Bo Wang, Peng Jia, Stephen J. Bush, Xia Wang, Yi Yang, Yu Zhang, Shijie Wan, Xiaofei Yang, Pengyu Zhang, Yuanting Zheng, Leming Shi, Lianhua Dong, Kai Ye

## Abstract

Recent advances in sequencing technologies have enabled the complete assembly of human genomes from telomere to telomere (T2T), resolving previously inaccessible regions such as centromeres and segmental duplications. Here, we present an updated, higher-quality, haplotype-phased T2T assembly of the Chinese Quartet (T2T-CQ), a family cohort comprising monozygotic twins and their parents, generated using high-coverage ONT ultralong and PacBio HiFi sequencing. The T2T-CQ assembly serves as a crucial reference genome for integrating publicly available multi-omics data and advances the utility of the Quartet reference materials. The T2T-CQ assembly scores highly on multiple metrics of continuity and completeness, with Genome Continuity Inspector (GCI) scores of 77.76 (maternal) and 76.41 (paternal), quality values (QV) > 70, and Clipping Reveals Assembly Quality (CRAQ) scores > 99.6 for both haplotypes, enabling complete annotation of centromeric regions. Within these regions, we identified novel 13-mer higher-order repeat patterns on chromosome 17 which exhibited a monophyletic origin and emerged approximately 230 thousand years ago. Overall, this work establishes an essential genomic resource for the Han Chinese population and advances the development of a T2T pan-Chinese reference genome, which will significantly enable future investigations both into population-specific structural variants and the evolutionary dynamics of centromeres.

## Introduction

The completion of the first telomere-to-telomere (T2T) human genome assembly, the primarily European T2T-CHM13 [1], was a landmark achievement in genomics, providing an updated reference for exploring the full complexity of the human genome, including previously unresolved regions such as centromeres, telomeres, rDNAs, and segmental duplications. This breakthrough was facilitated by advancements in long-read sequencing technologies, specifically Oxford Nanopore Technologies (ONT) and PacBio high-fidelity (HiFi) platforms, which enabled comprehensive resolution of highly repetitive and structurally complex genomic regions, contributing to a more complete understanding of human genetic variation [2–4]. Several high-quality Chinese reference genomes, including YH2.0 [5], HX1 [6], NH1.0 [7], Han1 [8], and HJ [9], have also been successfully assembled and released. However, these assemblies remain either partially collapsed or contain unresolved gaps, indicating room for further improvement in completeness and accuracy. Following this, and to close gaps in ethnic diversity, the T2T-CN1 [10] and T2T-Yao [11] assemblies were released, representing complete diploid genomes of Han Chinese individuals and highlighting the importance of high-quality, population-specific reference genomes to capture the full spectrum of complex genetic variation [12]. Building on these achievements, we present the T2T-CQ genome, a high-quality, haplotype-resolved T2T assembly generated from a family of Han Chinese descent [13, 14]. The Chinese Quartet project (https://chinese-quartet.org/) is a suite of DNA, RNA, protein and metabolite reference materials from B-lymphoblastoid cell lines (LCLs) from two monozygotic twin daughters (LCL5 and LCL6) and their biological parents (LCL7 and LCL8 for father and mother, respectively), extensively detailed in previous publications [13–18]. Our initial assembly of the CQ family [16], hereafter CQ v2.0, laid the groundwork for the present study in which we refine and expand it. By incorporating high-depth ultralong ( > 100 kb) ONT reads we significantly enhance both the contiguity and completeness of CQ v2.0 to T2T-level quality, with both maternal and paternal haplotypes attaining average quality values over 70.

Centromeres, which are essential for chromosome segregation during cell division, remain among the most challenging regions to assemble due to their highly repetitive nature and structural complexity [2, 19]. T2T-level assemblies therefore enable comprehensive annotations of centromeric regions, including the identification of higher-order repeats (HORs), which are essential for understanding centromere structure and function [3, 20, 21]. As such, we focus on characterizing centromeric regions in the T2T-CQ genome, identifying novel HORs and investigating their emergence and evolution. Additionally, we evaluate the performance of two state-of-the-art genome assembly tools, hifiasm and verkko, using our twin sequencing data as a robust benchmark. With the availability of a T2T-level reference genome, the Chinese Quartet provides an invaluable model for studying centromere biology, genetic variation, and evolutionary dynamics.

## Results and Discussion

### A diploid T2T genome assembly for the Chinese Quartet

In previous work [16], we presented CQ v2.0, haplotype-resolved assemblies of the monozygotic twins. However, although these assemblies were highly contiguous, they fell short of T2T standard, with the maternal haplotype assembly (CQ v2.0_mat) containing 34 gaps, and the paternal (CQ v2.0_pat) 51 gaps (**Table 1**). Moreover, 17 and 20 telomere motifs were missing in the maternal and paternal CQ v2.0 assemblies, respectively (Table 1). In this study, we supplemented our previously used set of 10.3× ONT ultralong reads ( > 100 kb) with an additional 57.3× (Table S1), aiming to assemble a T2T-level haplotype-resolved genome to serve as a new reference genome for the CQ. We constructed the initial assemblies using two state-of-the-art tools, verkko [22, 23] and hifiasm [24], in trio binning mode. As described in the methods and illustrated in (Figure S1A), the contigs from the two assembly results were merged for the maternal haplotype, leaving 9 gaps distributed across chromosomes 1, 6, 10, 11, 13, 14, 15, 21, and 22. Except for the short arms of acrocentric chromosomes (SAACs) 13, 14, 15, 21, and 22, gaps in the other chromosomes were filled using TGS-GapCloser [25] in conjunction with binning ONT reads and the ONT-based hifiasm assembly (Figure S1B). For the paternal haplotype, 8 gaps remained, located on chromosomes 4, 5, 13, 14, 15, 21, 22, and X, which we also filled using TGS-GapCloser. The rDNA clusters in the SAACs of both maternal and paternal haplotypes were finalized by adopting the strategy employed for T2T-CN1 [10]. Briefly, this involved predicting the rDNA copy number (CN) using droplet digital PCR (ddPCR), followed by evaluating the rDNA copy number for each chromosome using the T2T-CHM13 major rDNA morphs [26]. The ddPCR results show that, as expected, the copy ratios of the two twins were similar, with values of 276.92 and 283.73, respectively. Since we combined data from the twins (see Materials and Methods), we used their average value of 280 as the rDNA CN. Following the approach described in the “rDNA analysis” section of the methods, we found that the rDNA CNs for CQ v3.2_mat and CQ v3.2_pat were 123 and 157, respectively (Table 1). The five rDNA major morphs used for genome filling correspond to the five SAACs, with the rDNA CN for each of these chromosomes in CQ v3.2_mat and CQ v3.2_pat detailed in Table S2 and Table S3, respectively. We polished the assembled genome using NextPolish2 [27], ultimately obtaining T2T diploid human genome assemblies, designated as CQ v3.0_mat (maternal haplotype) and CQ v3.0_pat (paternal haplotype) (**Figure 1**A). Subsequently, we integrated the mitochondrial genome into the maternal haplotype assembly and performed structural variant (SV) error correction using a previously published pipeline [28], followed by manual validation. This refinement process identified and resolved three SVs in CQ v3.0_pat (Figure S2), whereas no SVs were detected in CQ v3.0_mat. The resulting high-quality T2T assembly was designated as the final CQ v3.2 genome. These two haplotype-resolved assemblies were further validated using high-throughput chromosome conformation capture (Hi-C) contact maps and manual inspection, which confirmed the absence of significant assembly errors (Figure S3A,B).

**Figure 1.**
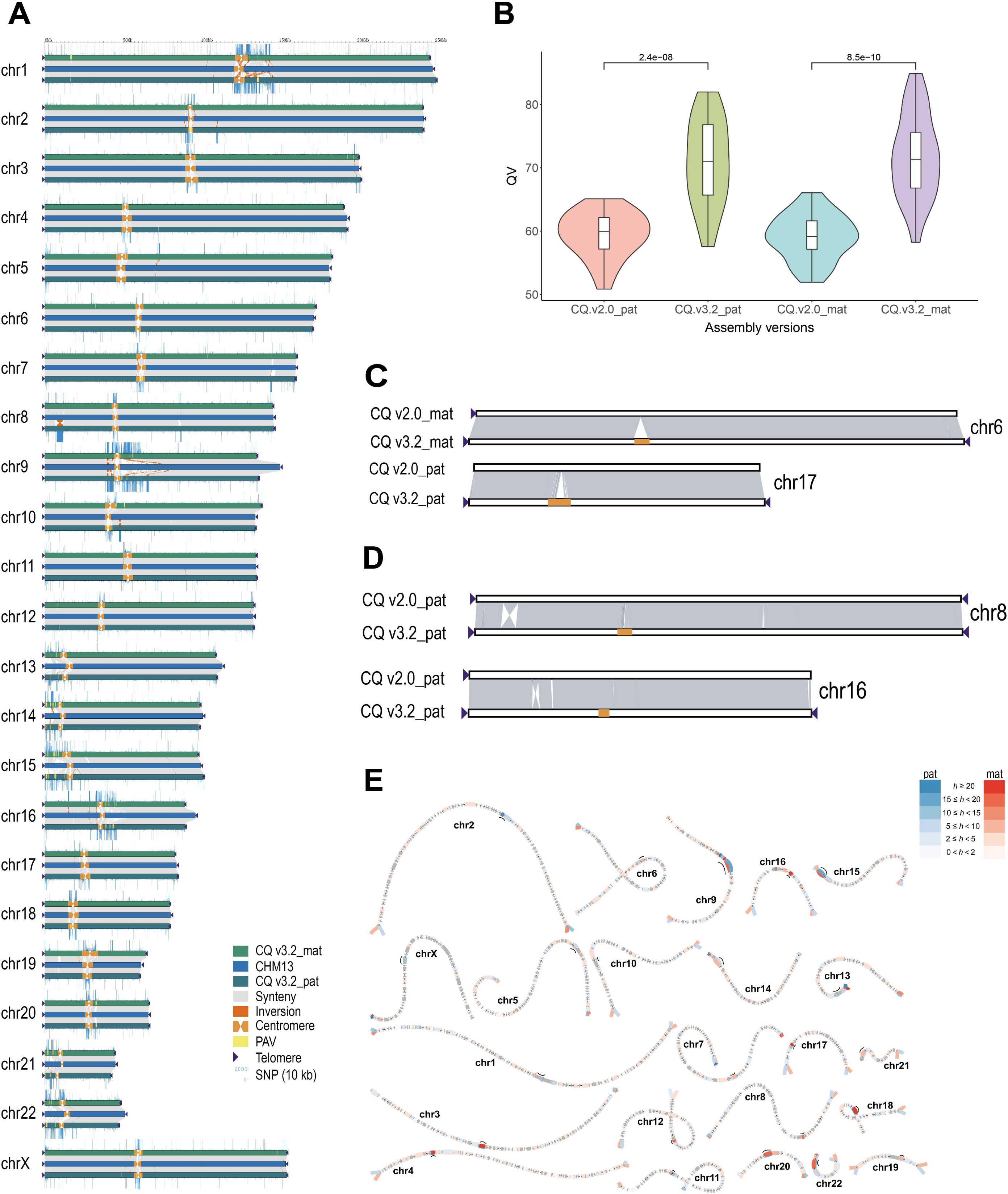
Overview of CQ v3.2 and its comparison with CQ v2.0. (**A**) Comparison of the CQ v3.2 and CHM13 reference genomes. Each set of chromosomes includes the maternal haplotype of CQ v3.2 (CQ v3.2_mat, top), the paternal haplotype of CQ v3.2 (CQ v3.2_pat, bottom), and CHM13 (middle). (**B**) Comparison of the QV (k = 21) for each chromosome of the maternal and paternal haplotypes in CQ v3.2 and CQ v2.0. (**C**) Illustration of the advantage of CQ v3.2 over CQ v2.0 in centromeric region assembly. (**D**) Illustration of large inversion variation between CQ v3.2 and CQ v2.0. (**E**) Visualization of heterozygous regions between two haplotypes using bubbles. Regions with different heterozygous variants are displayed in distinct colors, while centromeric regions are represented by black lines. The heterozygosity rate (*h*) is calculated as the structural variant count in each 500 kb window.

**Table 1.** Comparison of assembly quality among human telomere-to-telomere genome assemblies.

| Statistic | CQ.v2.<br>0_mat | CQ.v2.<br>0_pat | CQ.v3.<br>2_mat | CQ.v3.<br>2_pat | CN1.v1.<br>0.1_mat | CN1.v1.<br>0.1_pat | YAOv1.<br>1_mat | YAOv1.<br>1_pat |
| --- | --- | --- | --- | --- | --- | --- | --- | --- |
| Assembly size (Gb) | 3.00 | 3.00 | 3.03 | 3.03 | 3.04 | 2.94 | 3.02 | 2.92 |
| No. of gaps | 34 | 51 | 0 | 0 | 0 | 0 | 0 | 0 |
| No. of telomeres | 29 | 26 | 46 | 46 | 46 | 46 | 46 | 46 |
| Contig N50 (Mb) | 132.84 | 132.84 | 155.11 | 155.17 | 157.36 | 145.78 | 155.06 | 145.07 |
| QV range among chromosomes (k = 21) | 51.93-66.05 | 50.85-65.11 | 58.25-84.87 | 57.57-81.97 | 47.33-70.83 | 51.91-73.02 | 60.58-85.47 | 63.39-91.52 |
| QV average of chromosomes (k = 21) | 59.07 | 59.43 | 71.72 | 70.82 | 64.98 | 63.17 | 72.23 | 74.47 |
| QV whole genome (k = 21) | 57.55 | 58.29 | 67.90 | 66.65 | 60.11 | 59.37 | 70.49 | 72.28 |
| QV range among chromosomes (k = 31) | 49.59-64.75 | 50.65-63.73 | 60.45-76.80 | 59.50-80.23 | NA | NA | NA | NA |
| QV average of chromosomes (k = 31) | 57.19 | 57.50 | 67.94 | 68.43 | NA | NA | NA | NA |
| QV whole genome (k = 31) | 55.70 | 56.65 | 66.29 | 65.94 | NA | NA | NA | NA |
| False duplication (%) (k = 21) | 0.37 | 0.40 | 0.12 | 0.10 | NA | NA | 0.30 | 0.20 |
| False duplication (%) (k = 31) | 0.46 | 0.49 | 0.19 | 0.17 | NA | NA | NA | NA |
| Haplotype switch error (%) (k = 21) | 0.16 | 0.14 | 0.19 | 0.17 | NA | NA | 0.02 | 0.01 |
| Haplotype switch error (%) (k = 31) | 0.25 | 0.22 | 0.33 | 0.27 | NA | NA | NA | NA |
| Completeness (%; haploid) (k = 21) | 97.87 | 97.79 | 97.92 | 97.92 | NA | NA | 99.65 | 99.59 |
| Completeness (%; haploid) (k = 31) | 96.91 | 97.00 | 97.10 | 97.10 | NA | NA | NA | NA |
| Completeness (%; diploid) (k = 21) | 99.51 |  | 99.54 |  | NA | NA | NA | NA |
| Completeness (%; diploid) (k = 31) | 99.52 |  | 99.61 |  | NA | NA | NA | NA |
| GCI (HiFi + ONT) | NA | NA | 77.76 | 76.41 | 66.79 | 77.90 | NA | NA |
| Chromosome composition | 22 + X | 22 + X | 22 + X | 22 + X | 22 + X | 22 + Y | 22 + X | 22 + Y |
| rDNA copy number | NA | NA | 123 | 157 | 132 | 117 | 79 | 149 |

**Note:** Merquy results of CQ.v2.0 and CQ.v3.2 from the same pipeline used in this study. The QV and GCI data for CN1 were obtained from [10] and [33], respectively; the data for YAO were obtained from [11]. NA, not available.

In addition, we evaluated CQ v3.2 using Merqury [29], a k-mer-based reference-free assessment pipeline, using both PCR-free pair-end and HiFi reads to identify assembly errors (Figure S4A) and switching errors between the CQ v3.2_mat and CQ v3.2_pat haplotypes (Table 1; Figure S4B). We observed a higher switch error rate in CQ v3.2 compared to CQ v2.0, consistent across both 21-mer and 31-mer databases (Table 1). This can be attributed to the fact that CQ v2.0 was assembled by phasing long reads using the read-based phasing tool WhatsHap [30] followed by separate haplotype assembly [16]. While this method reduces switch errors, it compromises assembly continuity [16]. In contrast, CQ v3.2 adopted a trio binning approach instead, which prioritizes continuity at the cost of occasional switch errors [31]. Notably, despite this slight increase, CQ v3.2’s switch error rates (0.19% and 0.17% for 21-mers; 0.33% and 0.27% for 31-mers) remain substantially lower than the average haplotype switch error rate (0.67% for 21-mers) reported among 47 high-quality diploid human genome assemblies [32]. As such, CQ v3.2 demonstrates superior completeness compared to CQ v2.0 while exhibiting significantly lower false duplication rates (Table 1). Notably, CQ v3.2’s false duplication rate is even lower than that of T2T-Yao (Table 1).

The chromosome-wide average quality values (QV) (k = 21) for CQ v3.2_mat (71.72) and CQ v3.2_pat (70.82) demonstrate significant improvement over both CQ v2.0_mat (59.07) and CQ v2.0_pat (59.43) (Figure 1B), substantially exceeding those of the first T2T diploid human genome assembly, CN1 [10] (Table 1). The QV values of CQ v3.2 were nearly comparable to those of T2T-Yao [11] and approached the quality of T2T-CHM13 (QV = 73.94) in a parallel comparison [28]. We additionally performed Merqury QV assessments using 31-mers to compare CQ v3.2 and CQ v2.0. Due to the increased sensitivity of 31-mers, particularly in homopolymer-rich regions, the QV scores (k = 31) were generally lower than those obtained with 21-mers (Table 1). Nevertheless, CQ v3.2 consistently maintained higher QV values than CQ v2.0 (Table 1).

Moreover, compared to CQ v2.0, the contig N50 and QV of CQ v3.2 were significantly improved (Table 1). The contig N50 values in CQ v3.2_mat and CQ v3.2_pat are 155.11 Mb and 155.17 Mb, respectively, surpassing those of CQ v2.0_mat (132.84 Mb) and CQ v2.0_pat (132.84 Mb). To further evaluate the assembly continuity of CQ v3.2, we employed two computational tools: Genome Continuity Inspector (GCI) [33] and Clipping Reveals Assembly Quality (CRAQ) [34]. Our CQ v3.2 assemblies achieved GCI scores (HiFi+ONT) of 77.76 (CQ v3.2_mat) and 76.41 (CQ v3.2_pat), demonstrating continuity comparable to T2T-CN1_pat (77.90) while showing substantial improvement over T2T-CN1_mat (66.79) (Table 1). Using CRAQ, we obtained both regional assembly quality indicators (R-AQI) and structural assembly quality indicators (S-AQI), with CQ v3.2_mat scoring R-AQI = 99.61 and S-AQI = 99.82, while CQ v3.2_pat achieved R-AQI = 99.65 and S-AQI = 99.79. According to CRAQ’s quality thresholds, an AQI score greater than 90 indicates reference-level quality [34]. We evaluated the CQ v3.2 genomes using HMM-Flagger [32], a read-mapping-based tool designed to detect mis-assemblies in diploid genomes. Based on HiFi and ONT ultralong read mappings, 99.33% and 99.17% of the CQ v3.2 assembly, respectively, was flagged as correctly assembled (Table S4). Overall, the convergence of multiple evaluation metrics provides robust evidence for the exceptional continuity and accuracy of our CQ v3.2 assemblies compared to CQ v2.0, particularly in centromeric regions (Figure 1C). A representative example is observed on chromosome 6: while the CQ v2.0_mat assembly spans 167,988,758 bp, the CQ v3.2_mat assembly extends to 173,304,238 bp. This represents a gain of 5,315,481 bp of centromeric sequence in the T2T-level assembly (Figure 1C; Table S5).

Comparative analysis of the CQ v3.2 and CQ v2.0 genomes also identified two megabase-scale inversions on chromosomes 8 and 16 (Figure 1D, Figures S5 and S6), specifically within the paternal haplotypes. These structural variants were validated through GCI-based sequencing coverage plots (Figure S7) and manual verification (Figure S8), confirming their accuracy. We next used SyRI [35] to identify all inversion variants > 100 kb between CQ v3.2 and CQ v2.0. A total of 7 and 13 inversions were detected in the maternal and paternal haplotypes, respectively (Table S6), which we again confirmed with GCI and manual verification (Figures S7–S10).

The heterozygosity rate between the two CQ v3.2 haplotype genomes, calculated using the “SV count” method (https://github.com/T2T-CN1/CN1/tree/main/heterozygosity), revealed that the most diverse regions are located in the centromeres (Figure 1E). The heterozygosity rate in the centromeric regions is 5.3 times higher than that in the non-centromeric regions (on average 12.18 vs. 2.29 SVs per 500 kb) (Figure S11), similar to the 5.4-fold difference observed in T2T-CN1 [10]. The high heterozygosity in centromeric regions suggests that these areas are genetically diverse and may play a significant role in maintaining genetic variation within a population.

Since our genome assembly was constructed by merging data from monozygotic twins LCL5 and LCL6, we identified variants in each twin using CQ v3.2 as reference to enrich the heterozygous sites in the assembly. Our analysis detected 3,663 SNVs and 20,680 indels (Table S7, available at https://github.com/BoWangXJTU/T2T-CQ/tree/main/twin_diff), including 1027 SNVs and 109 indels with allele frequencies > 0.3 located outside repeat regions (Figures S12 and S13; Table S7).

Notably, we found that the maternal haplotype contained 142 SNVs and 34 indels unique to LCL5, compared to 314 SNVs and 50 indels unique to LCL6 (Table S7). Similarly, for the paternal haplotype, 145 SNVs and 37 indels were specific to LCL5, while 426 SNVs and 68 indels were unique to LCL6 (Table S7). These comprehensive variant profiles provide a valuable resource for future studies in human family genetics [4].

### Centromeric annotations and evolutionary dynamics of novel HORs

With the generation of T2T-CQ, we were able to characterize the landscape of centromeric sequences in the Chinese Quartet. First, we annotated the centromeric α-satellite regions in CQ3.2_mat and CQ3.2_pat using the CenMAP pipeline, which ranged from 2.7 Mb to 10.2 Mb in CQ3.2_mat (Table S8) and from 2.5 Mb to 8.4 Mb in CQ3.2_pat (Table S9). Sequence identity maps generated using StainedGlass [36] are displayed for each centromere in both CQ3.2_mat and CQ3.2_pat (Figures S14–S17). The length of the α-satellite HOR array is well known to exhibit variation across human populations [3, 37]. Accordingly, we annotated the HOR array lengths in both haplotypes and compared them with the HOR lengths of 102 additional human samples [37] (detailed in Materials and Methods) (**Figure 2**A). We found that the HOR length of CQ3.2_mat ranged from 787 kb to 6.3 Mb (Table S10), while that of CQ3.2_pat ranged from 183 kb to 5.1 Mb (Table S11). Interestingly, the HOR lengths of the two haplotypes of CQ differ significantly, with the exception of chromosome 2, where the HOR lengths are similar (1,999,710 bp in CQ3.2_mat and 1,978,117 bp in CQ3.2_pat), both close to the mean value (Figure 2A).

**Figure 2.**
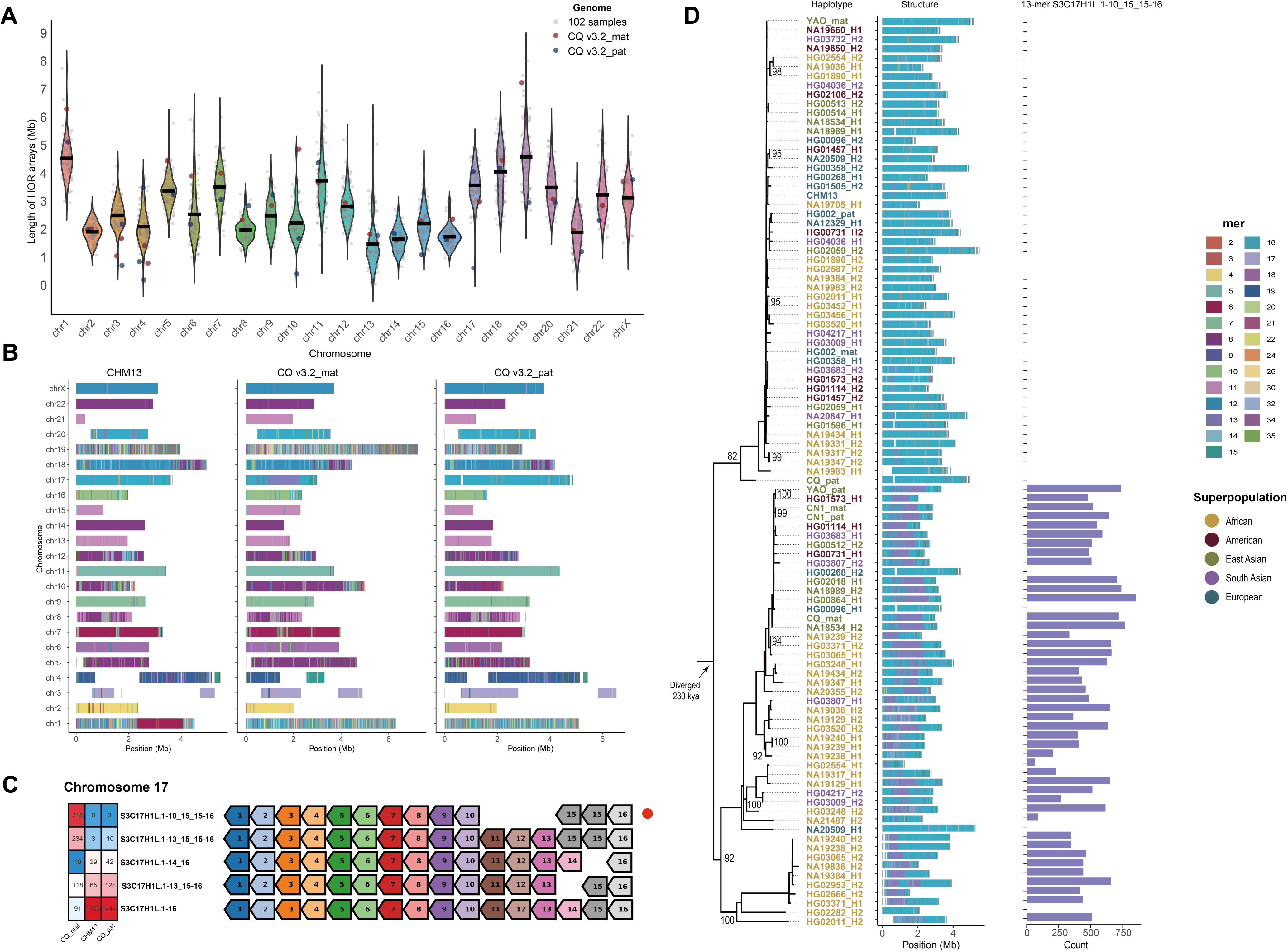
Comparative analysis of centromeric higher-order repeat (HOR) length, structural organization, and evolutionary emergence of novel HORs in CQ v3.2 versus diverse human genomes. (**A**) The length of the HOR arrays in CQ v3.2_mat (red), CQ v3.2_pat (purple), and 102 human samples. (**B**) The HOR array structure is shown on the axes, along with the organization of each centromeric region in CHM13, CQ v3.2_mat, and CQ v3.2_pat. (**C**) The heatmap showing the HOR number in chromosome 17 of CHM13, CQ v3.2_mat, and CQ v3.2_pat, along with the HOR patterns. Monomer composition is indicated by numbered arrows (color-coded by type), with novel HOR patterns marked by red solid circles. (**D**) Maximum-likelihood phylogeny of chromosome 17 HOR arrays, with divergence time estimates supporting a monophyletic origin for novel 13-mer HORs. Nodes with ≥ 80% bootstrap support are labeled numerically. Bar plots quantify 13-mer HOR abundance across five superpopulations (African, American, East Asian, South Asian, European), with haplotype colors corresponding to their geographic origin.

We subsequently visualized and compared the α-satellite HOR array structures between the CQ v3.2 haplotypes (Chinese) and T2T-CHM13 (European) (Figure 2B), finding substantial differences in HOR composition between them. To assess population-level variation, we systematically analyzed divergence patterns across centromeric contigs, including both European-Chinese comparisons and intra-Chinese individual variations (see Materials and Methods). Notably, no significant correlation was detected between intra- and inter-population variation patterns (Figures S18 and S19), which may be explained by the dynamic evolutionary behavior of HORs [3].

For chromosome 1, the T2T-CHM13 assembly exhibits nearly 2 Mb of unique HOR structures compared to both CQ3.2_mat and CQ3.2_pat (Figure 2B). These regions predominantly comprise 2-mer HORs (S1C1/5/19H1L.5_6/4 and S1C1/5/19H1L.5-6) and a 6-mer HOR (S1C1/5/19H1L.1-5_6/4). Strikingly, the S1C1/5/19H1L.5_6/4 motif appears to be entirely absent from chromosome 1 in both haplotypes of CQ v3.2, CN1, YAO_mat, and HG002 (Table S12). Interestingly, YAO_pat shows a HOR structure remarkably similar to CHM13 (Figure S20), suggesting that these two haplotypes may have independently acquired and expanded hybrid monomers. This pattern parallels the observed hybridization between S1C1/5/19H1L monomer 6 and monomer 4.

For chromosome 17, a unique region of less than 2 Mb is observed in CQ v3.2_mat, composed of the 13-mer HOR S3C17H1L.1-10_15_15-16 (Figure 2B). The canonical HOR pattern for chromosome 17 is S3C17H1L.1-16 [10], whereas S3C17H1L.1-10_15_15-16 lacks monomers 11, 12, 13, and 14, and has monomer 15 duplicated once (Figure 2C). The S3C17H1L.1-10_15_15-16 motif appears exclusively in CQ v3.2_mat, while it is entirely absent in T2T-CHM13 and present in only three copies in CQ v3.2_pat (Figure 2C). This suggests that S3C17H1L.1-10_15_15-16 represents a novel HOR specific to CQ v3.2 (Figure 2B,C). Interestingly, both alleles of chromosome 17 in T2T-CN1 and one haplotype in T2T-YAO were found to contain this 13-mer HOR [10] (Figure S21), initially suggesting a potential association with the Han Chinese population.

To test this hypothesis, we analyzed 64 additional genomes from HGSVC3, HG002, and T2T-YAO, representing diverse superpopulations (African, American, European, South Asian, and other East Asian groups). Among 99 fully assembled chromosome 17 centromeres subjected to HOR annotation, 43 haplotypes (43/99) carried the S3C17H1L.1-10_15_15-16 motif across all superpopulations (Figure 2D), indicating that this “novel 13-mer HOR” is not unique to Han Chinese. To investigate the evolutionary dynamics underlying these HOR arrays – particularly the emergence of novel HORs – we constructed phylogenetic trees using chimpanzee sequences as an outgroup and estimated divergence times, assuming a human-chimpanzee split at 6 million years ago (Mya) (Figure 2D). Phylogenetic analysis revealed two distinct clusters, with haplotypes containing the novel HORs predominantly forming a monophyletic clade (Figure 2D). We estimate that these novel HORs of chromosome 17 arose approximately 230 thousand years ago (kya). While similar HOR evolutionary patterns were reported for chromosomes 5, 10, 11, and 12 [3], our findings on chromosome 17 uncover a more widespread distribution of novel HORs across diverse populations, suggesting additional layers of centromeric evolution.

### Twin-based validation of centromeric assembly fidelity

Our CQ material differs from the CN1 and YAO assemblies in comprising sequencing data from two monozygotic twins, representing two distinct trios at the data level. This makes them particularly suitable for evaluating trio binning assembly software, especially in complex centromeric regions. In our systematic evaluation using ONT ultralong coverages (ranging from 5× to a maximum coverage of 40.4× for LCL5 and 27.2× for LCL6, with HiFi reads normalized to 50× for both samples), we employed two state-of-the-art assemblers (hifiasm v0.25.0 and verkko v2.2.1) to assess centromeric region assembly robustness. The results demonstrated that both assemblers generally produced improved contig N50 values with increasing ONT ultralong coverage, enabling more complete centromere assembly in both twins. Notably, while LCL5 assemblies in hifiasm showed a slight N50 decrease beyond 20× ONT coverage (at 25× and 40.4×), they still yielded more complete centromeres than at 20× (Table S13).

We next performed comparative analyses of complete centromere lengths across different coverage depths (Figure S22A,B), revealing both assemblers to be comparably robust. Further detailed analysis of maximum length variations for each centromere at ≥ 15× sequencing depth showed that hifiasm assemblies exhibited a maximum length difference of 16,629 bp in LCL5 (chromosome 10 paternal haplotype) and 182,966 bp in LCL6 (chromosome 15 paternal haplotype) (Table S14). Similarly, verkko assemblies displayed maximum differences of 28,880 bp in LCL5 (chromosome 7 maternal haplotype) and 75,499 bp in LCL6 (chromosome 11 paternal haplotype).

These coverage-dependent variations were significantly smaller than allelic differences. For instance, in LCL6’s chromosome 15, while maternal/paternal haplotype differences reached 1.3 Mb, the maximum length variations within maternal (37,284 bp) and paternal (8019 bp) haplotypes were orders of magnitude smaller.

Our findings demonstrate that both assemblers can control centromeric assembly errors within 300 kb (often to just a few bp), minimizing impact on sample haplotype comparisons. However, for base-level or HOR-level analyses, further improvements in assembly precision remain necessary.

## Conclusions

In summary, this study significantly advances genomic research by presenting a high-quality T2T diploid assembly of the Chinese Quartet. Leveraging cutting-edge sequencing technologies, we identified novel HORs in centromeric regions and elucidated their dynamic evolutionary patterns, including expansions and contractions. Additionally, we systematically evaluated the robustness of state-of-the-art assembly tools in centromere reconstruction using twin sequencing data. These findings deepen our understanding of centromere structural diversity and evolution. Moreover, this work establishes a critical genomic resource for constructing a comprehensive T2T pan-Chinese reference genome, which will facilitate population-scale genomics research and precision medicine initiatives. [38, 39].

## Materials and Methods

### Sample collection

The Chinese Quartet family, comprising a 60-year-old father (LCL7), 60-year-old mother (LCL8), and their two 30-year-old monozygotic twin daughters (LCL5 and LCL6), all from the Fudan Taizhou cohort, are Certified Reference Materials according to the State Administration for Market Regulation in China (reference numbers GBW09900-GBW09903). The process of establishing the cell lines was described in a prior study [40].

### ONT ultralong library preparation and sequencing

ONT ultralong DNA was extracted using the GrandOmics Genomic DNA Kit following the manufacturer’s guidelines. The quality and quantity of the total DNA were assessed to ensure suitability for ultra-long Nanopore library construction, which was performed according to the established protocols [41]. DNA libraries (approximately 400 ng) were constructed and sequenced on the PromethION platform (Oxford Nanopore Technologies) at the Genome Center of GrandOmics (Wuhan, China).

### Hi-C sequencing

The Hi-C library was prepared from cross-linked chromatins of the cells following a standard Hi-C protocol [42]. The library was subsequently sequenced on the MGI-2000 platform. The Hi-C sequencing data were used to map the assembled genome using Juicer v1.6 [43]. The assemblies were then manually reviewed and visualized using Juicebox v1.11.08 [44].

### Genome assembly

Since monozygotic twins are generally considered genetically identical, with only limited somatic substitutions [45], we merged the sequencing reads from the two children to generate a high-quality haplotype-resolved genome [16]. The initial assemblies were constructed using state-of-the-art tools, Verkko v2.2.1 [22, 23] and hifiasm v0.19.9 [24], in trio mode. The scaffolding of the Verkko assemblies was enhanced using the hifiasm assembly, as the contig N50 of the Verkko-assembled genome (131.8 Mb) was greater than that of hifiasm (107.5 Mb). For each haplotype, hifiasm and Verkko assemblies were assigned to their respective chromosomes and aligned to the CHM13 reference genome (Figure S1A). Within each chromosome, contigs were merged into longer scaffolds based on overlaps between hifiasm and Verkko assemblies. Additionally, supplementary assemblies were generated using hifiasm v0.24.0 with the ‘--ONT’ parameter, incorporating canu-based [46] binned ONT reads. The gaps were initially filled using TGS-GapCloser v 1.2.1 [25], leveraging the hifiasm assemblies with ONT data and binned ONT reads. The highly repetitive nature of rDNA, combined with heterozygosity, presents significant challenges for resolution [1]. Therefore, only the rDNA copy number was estimated to populate each chromosome with corresponding identical copies [10]. The mitochondrial genome was assembled using the software Unicycler v0.5.1 [47].

### Genome polishing

Since our genome assembly was performed using the trio binning approach, the two haplotype assembly results were polished separately. First, the yak v0.1-r56 (https://github.com/lh3/yak) was used to bin HiFi reads based on parental information. Subsequently, the ‘getseq’ function from fxTools v0.1.0 (https://github.com/moold/fxTools) was employed to extract the binned HiFi reads. The “Racon + Merfin” pipeline was then executed to generate the HiFi mapping file [28], “racon.meryl.iter_1.winnowmap.bam”. Finally, the NextPolish2 workflow was run to produce the polished genome files [27]. To identify and correct potential SV errors, we implemented the “Find SV-like errors” step from the T2T polishing pipeline (https://github.com/arangrhie/T2T-Polish) to refine our genome assembly. [28].

### Genome assessment

To construct a hybrid k-mer database, we employed Meryl v1.4.1 to extract 21-mers and 31-mers from both Illumina PCR-free paired-end data and HiFi reads obtained from the twins (Table S1). The database was generated using the following sequence of commands:

meryl greater-than 1 twins.hifi.meryl output twins.hifi.gt1.meryl

meryl greater-than 1 twins.k31mer.meryl output twins.k31mer.gt1.meryl

meryl divide-round 3 twins.k31mer.gt1.meryl output twins.k31mer.gt1.divideRound3.meryl

meryl union-max twins.hifi.gt1.meryl twins.k31mer.gt1.divideRound3.meryl output hybrid.meryl

Parental datasets, consisting of Illumina PCR-free paired-end sequences, were obtained from LCL7 (father) and LCL8 (mother) (Table S1). Subsequently, Merqury was employed to evaluate the assemblies [29], following the methodology outlined in previous studies [10, 11]. In addition to k-mer-based metrics, assembly continuity was assessed using GCI v1.0 [33] and CRAQ v1.0.9 [34], two read-mapping-based tools that incorporate both HiFi and ONT ultra-long reads. We subsequently employed the HMM-Flagger v1.1.0 tool to identify erroneous regions, false duplications, and collapsed segments in the diploid genome assembly [48], while also detecting structurally correct haploid regions. HMM-Flagger analysis was performed separately using read depth information, HiFi reads, and ONT ultralong reads.

### Genomic variation

To comprehensively identify the full spectrum of heterozygosity between the CHM13 and the diploid T2T-CQ genomes, we performed a direct comparison of these three assemblies using minimap2 v2.20-r1061 [49] with the following parameters: -x asm5 -t 64 --cs. The paf2delta.pl script (https://github.com/gorliver/paf2delta) was used to convert the PAF format into the delta format. Subsequently, the ‘show-snps’ function from MUMmer v4.0.0rc1 [50] and the ‘Transform --SNP’ function from GenomeSyn [51] were employed to generate SNP results. Additionally, the ‘show-diff’ function from MUMmer and the ‘Transform --PAV’ function from GenomeSyn were utilized to produce PAV (Presence-Absence Variation) results. Following this, GenomeSyn was utilized to conduct a whole-genome comparison between the CHM13 genome and the assembled genome in this study. Structural variants > 100 kb, specifically inversions, between the CQ v3.2 and CQ v2.0 genome assemblies were identified using SyRI v1.7.0 [35]. For heterozygous sites comparison in twins, phased HiFi reads from both twins were aligned to the maternal and paternal haplotypes of the CQ v3.2 genome using “minimap2 + VarScan v2.4.6” variant calling pipeline [49, 52].

### Centromere analysis

CenMAP v0.2.4 (https://github.com/logsdon-lab/CenMAP), a centromere mapping and annotation pipeline designed for T2T human genome assemblies, was utilized to predict potential centromeric regions in the genome. The α-satellite sequences were annotated using the pipeline hmmer-run.sh (https://github.com/fedorrik/HumAS-HMMER_for_AnVIL) with the hmm file ‘AS-HORs-hmmer3.3.2-120124.hmm’. Subsequently, HOR structural variants were identified using the tool StV (https://github.com/fedorrik/stv). StainedGlass v0.6 was employed to visualize the centromeric regions [36], and custom R scripts were utilized to plot the HOR length and StV annotations. The HOR length was calculated using ‘censtats’ from CenMAP v0.2.4. The HOR lengths of 102 additional human samples – including 22 African from Human Pangenome Reference Consortium (HPRC) [32], 16 Latin American (HPRC), 62 East Asian (4 from HPRC and 58 from Chinese Pangenome Consortium [48]), and 1 South Asian (HPRC) individuals, plus CHM13 – were obtained from our previous study [37]. For comprehensive analysis of centromere structure and phylogeny, we integrated multiple recently available genomic resources, including 64 Human Genome Structural Variation Consortium Phase 3 (HGSVC3) genomes [12] (excluding HG002, for which we used v1.1 from https://github.com/marbl/HG002), YAOv1.1 [11], CN1v1.0.1 [10], CHM13 [1], and CQv3.2. In total, this dataset comprised 69 samples representing 137 haplotypes.

### Phylogenetic analysis

To investigate the phylogenetic relationships and divergence times of centromeric regions from chromosome 17 across 69 diverse human genomes (encompassing 137 haplotypes), we conducted phylogenetic analyses following a modified approach from [3]. We first identified contigs containing fully assembled centromeres using CenMAP v0.2.4, then aligned each to the unique q-arm reference region (chr17: 28,103,337–28,138,133, CHM13 v2.0) using unimap (https://github.com/lh3/unimap). For the outgroup comparison, we included chimpanzee sequences [53] aligned to the same unique region. Multiple sequence alignment was performed using MAFFT v7.526 [54] (with parameter --auto), followed by maximum-likelihood phylogenetic reconstruction with IQ-TREE v2.0.3 [55] (with parameter -m MFP -B 1000 --alrt 1000 -T AUTO). The resulting trees were visualized in FigTree v1.4.4 (http://tree.bio.ed.ac.uk/software/figtree/). Divergence time estimation was conducted using r8s v1.81 [56] with the penalized likelihood method and the truncated Newton algorithm, applying a molecular clock calibrated to a human-chimpanzee divergence time of 6 million years [3].

### Sequence identity analysis across centromeric regions

To assess sequence conservation and population divergence, we compared centromeric regions between European and Chinese haplotypes. For the Chinese population, we analyzed two haplotypes each from the CN1v1.0.1 [10], YAOv1.1 [11], and CQv3.2 assemblies, while for the European population, we examined CHM13v2.0 along with two haplotypes each from the HG002v1.1, HG00171 (HGSVC3), and HG00268 (HGSVC3) assemblies. The European and Chinese references were CHM13v2.0 and YAOv1.1.mat, respectively. Pairwise alignment of centromeric contigs was performed using minimap2 with optimized parameters (-I 15G -K 8G -ax asm20 --secondary = no --eqx -s 2500) [3], retaining only primary alignments (SAMtools flag 4). We then partitioned alignments into consecutive 10 kb non-overlapping windows relative to the reference genomes and calculated sequence identity per window as: (matches)/(matches + mismatches + insertions + deletions). This approach allowed systematic comparison of intra- and inter-population variation while accounting for structural differences in centromeric regions [3].

### Evaluation of assembly fidelity in centromeric regions

To evaluate the robustness of centromere assembly in two state-of-the-art genome assemblers (hifiasm v0.25.0 and verkko v2.2.1), we employed 50× HiFi reads as the baseline and tested with ONT ultralong reads ( > 100 kb) at varying coverages (5×, 10×, 15×, 20×, 25×, and maximum coverage). Random subsampling of sequencing reads was performed using Rasusa [57] with the following parameters: “reads --coverage specific coverage --genome-size 3Gb”. Both assemblers were run in trio binning mode using default parameters and were tested on twin samples LCL5 and LCL6. For centromere assessment, we identified complete centromeres using CenMAP v0.2.4 and compared their lengths to determine the robustness of each assembly approach.

### Estimation of rDNA copy number using ddPCR

The ddPCR experimental protocol followed a previously published method [10], with the modification that the single-copy gene *RPPH1* was used instead of the *TBP1* gene. The rDNA forward primer was 5’-AACGTGAGCTGGGTTTAG-3’, the rDNA reverse primer was 5’-CTCGTACTGAGCAGGATTAC-3’, and the rDNA probe was 5’-CACATCATCAGTAGGGT-3’. The forward primer, reverse primer, and probe for *RPPH1* were 5’-GAGGGAAGCTCATCAGTGG-3’, 5’-CCCTAGTCTCAGACCTTCC-3’, and 5’-CCACGAGCTGAGTGC-3’, respectively. The rDNA copy number was calculated as the ratio of rDNA copies/µL to *RPPH1* copies/µL.

### rDNA analysis

Ribotin was used to extract binned ONT reads containing rDNA sequences [26]. Subsequently, the extracted ONT reads were mapped to the major morphs of CHM13 (https://github.com/maickrau/ribotin_paper_experiments) using minimap2 v2.20-r1061 [49]. We calculated the ONT read counts and base counts for each major morph of CHM13, determined the rDNA copy number for each morph, and used this information to fill the rDNA regions in the genome assembly (https://github.com/BoWangXJTU/CQ_T2T/tree/main/04.rDNA_analysis).

## Ethical statement

The Quartet Project received approval from the Institutional Review Board of the School of Life Sciences at Fudan University (BE2050). The study was conducted in accordance with the principles outlined in the Declaration of Helsinki. Four healthy volunteers from a family quartet, participating in the Taizhou Longitudinal Study in Taizhou, Jiangsu, China, were enrolled. Peripheral blood samples were collected from these individuals to establish human immortalized B-lymphoblastoid cell lines. All four donors provided signed informed consent forms prior to participation.

## Data availability

The raw sequencing data of the Chinese Quartet have been deposited in the Genome Sequence Archive for Human [58] (GSA-Human) at the NGDC, BIG, CAS / CNCB, under accessions HRA010594. The CQ v3.2 genome sequences have been deposited in the Genome Warehouse [59] (GWH) at the NGDC, BIG, CAS / CNCB, under accessions GWHFQEY00000000.1 (https://ngdc.cncb.ac.cn/gwh/Assembly/reviewersPage/QTUAvfMmiHZltZaUxCcCWxFecuVWwiibmxDtFkihzBKMrGaXpMpRupbeYIvWIUxg) and GWHFQEX00000000.1 (https://ngdc.cncb.ac.cn/gwh/Assembly/reviewersPage/PUoxmconKbDSwirCKLxosGjJhJHDSEoqNnGPwwljFowZiUFivSRdMgcIsyyWPOXk).

## Code availability

The pipelines and code for genome assembly, polishing, centromere analysis, and rDNA analysis are available on GitHub: https://github.com/BoWangXJTU/T2T-CQ. The code has also been submitted to BioCode at the National Genomics Data Center (NGDC), Beijing Institute of Genomics (BIG), Chinese Academy of Sciences (CAS) / China National Center for Bioinformation (CNCB) (BioCode: BT007908), which is publicly accessible at https://ngdc.cncb.ac.cn/biocode/tools/BT007908.

## CRediT author statement

**Bo Wang:** Investigation, Writing – original draft, Methodology, Visualization, Funding acquisition. **Peng Jia:** Investigation, Methodology, Visualization, Writing – review & editing. **Stephen J. Bush:** Writing – review & editing. **Xia Wang:** Validation, Investigation. **Yi Yang:** Validation, Investigation. **Yu Zhang:** Validation, Investigation. **Shijie Wan:** Methodology, Visualization. **Xiaofei Yang:** Methodology, Funding acquisition. **Pengyu Zhang:** Methodology. **Yuanting Zheng:** Investigation, Data curation. **Leming Shi:** Investigation, Data curation. **Lianhua Dong:** Conceptualization, Supervision, Investigation, Data curation, Writing – review & editing, Resources, Funding acquisition. **Kai Ye:** Conceptualization, Supervision, Methodology, Writing – review & editing, Resources, Funding acquisition. All authors have read and approved the final manuscript.

## Acknowledgements

This study was supported by the National Key R&D Program of China (grant numbers 2023YFF0613300 and 2022YFC3400300), the National Natural Science Foundation of China (grant numbers 32200510, 32400509, 32125009, 32422019 and 62172325), the Natural Science Foundation of Shaanxi Province (2024JC-JCQN-28), the Fundamental Research Funds for the Central Universities (xzy012024088), and the Scientific Research Program of Shaanxi Provincial Department of Education (23JK0290).

## Supplementary Material

**Figure S1 Illustration of the process of merging hifiasm and Verkko assemblies, followed by subsequent gap-filling steps.**

(**A**) The hifiasm and Verkko assemblies for maternal chromosome 11 were aligned to CHM13 (middle) to organize and orient the contigs. The red block in CHM13 indicates the centromeric region. (**B**) The gap in the maternal chromosome 1 was closed using hifiasm assemblies in ONT mode with binned ONT ultralong reads.

**Figure S2 Comparative IGV screenshots demonstrating structural variant refinement through polishing.**

IGV, Integrative Genomics Viewer.

**Figure S3 The Hi-C contact maps of the CQ v3.2 maternal (A) and paternal (B) haplotypes.**

Hi-C, high-throughput chromosome conformation capture.

**Figure S4 The quality of the diploid CQ v3.2 assembly.**

(A) The spectrum plots of 31-mers in the assembly and reads for evaluating assembly errors. (B) Hap-mer blob plot of the CQ v3.2 assembly. mat, maternal haplotype. pat, paternal haplotype.

**Figure S5 Karyotype alignments of CQ v3.2 maternal (top) and CQ v2.0 maternal (bottom) haplotypes.**

Blue regions represent centromeric regions.

**Figure S6 Karyotype alignments of CQ v3.2 paternal (top) and CQ v2.0 paternal (bottom) haplotypes.**

Blue regions represent centromeric regions.

**Figure S7 GCI coverage plots validate inversions in CQ v3.2 assemblies differing from CQ v2.0.**

GCI, genome continuity inspector.

**Figure S8 IGV map of binned ONT reads mapped to CQ v3.2 and CQ v2.0.**

We manually confirmed that the large inversions between chr8 and chr16 of the paternal haplotypes in CQ v3.2 and CQ v2.0 were due to assembly errors in CQ v2.0, with CQ v3.2 being correct. IGV, Integrative Genomics Viewer.

**Figure S9 Four IGV maps of binned ONT reads aligned to the CQ v3.2 and CQ v2.0 maternal haplotypes, which demonstrate the correction of assembly errors in CQ v2.0, specifically for inversions greater than 100 kb.**

IGV, Integrative Genomics Viewer.

**Figure S10 Four IGV maps of binned ONT reads aligned to the CQ v3.2 and CQ v2.0 paternal haplotypes, which demonstrate the correction of assembly errors in CQ v2.0, specifically for inversions greater than 100 kb.**

IGV, Integrative Genomics Viewer.

**Figure S11 Heterozygosity comparison between centromeric and non-centromeric regions.**

The heterozygosity rate was calculated as the SV count in each 500 kb window. Red squares represent the mean values of the data. This comparison highlights the significantly higher heterozygosity in centromeric regions compared to non-centromeric regions.

**Figure S12 Comparison of heterozygous SNV and indel allele frequencies in twins.**

SNV, single nucleotide variants; indel, insertions/deletions; mat, maternal haplotype; pat, paternal haplotype.

**Figure S13 Comparison of heterozygous SNV and indel allele frequencies in twins.**

SNV, single nucleotide variants; indel, insertions/deletions; mat, maternal haplotype; pat, paternal haplotype.

**Figure S14 StainedGlass plots of the CEN regions of chromosomes 1 to 6 in CQ v3.2.**

CEN, centromeric

**Figure S15 StainedGlass plots of the CEN regions of chromosomes 7 to 12 in CQ v3.2.**

CEN, centromeric

**Figure S16 StainedGlass plots of the CEN regions of chromosomes 13 to 18 in CQ v3.2.**

CEN, centromeric

**Figure S17 StainedGlass plots of the CEN regions of chromosomes 19 to 22 and chromosome X in CQ v3.2.**

CEN, centromeric

**Figure S18 Scatter plots of centromere sequence identity between CHM13 (y-axis) and YAO_mat (x-axis) reference genomes compared to CHN and EUR population samples.**

CHN, Chinese; EUR, European.

**Figure S19 Centromere sequence identity patterns between CHN and EUR populations, including within-population comparisons.**

CHN, Chinese; EUR, European.

**Figure S20 Higher-order repeat array structures of the chromosome 1 centromeric region, compared among the CHM13, HG002, CN1, CQ, and YAO genomes.**

**Figure S21 Higher-order repeat array structures of the chromosome 17 centromeric region, compared among the CHM13, HG002, CN1, CQ, and YAO genomes.**

**Figure S22 Comparative analysis of complete centromere lengths at varying coverage depths in LCL5 and LCL6 using both hifiasm (A) and verkko (B) assemblers.**

**Table S1 Sequencing depth statistics used in this study.**

**Table S2 Predicted rDNA copy numbers for the five short arms of acrocentric chromosomes in the CQ v3.2 maternal haplotype.**

**Table S3 Predicted rDNA copy numbers for the five short arms of acrocentric chromosomes in the CQ v3.2 paternal haplotype.**

**Table S4 Performance assessment of HMM-flagger for CQ v3.2 diploid genome assemblies.**

**Table S5 Comparative chromosome lengths of CQ v3.2 and CQ v2.0 genome assemblies.**

**Table S6 Coordinates of inversion variants ( > 100 kb) identified between the CQ v2.0 and CQ v3.2 genomes.**

**Table S7 Heterozygous sites comparison in the twin cell lines LCL5 and LCL6.**

**Table S8 CQ v3.2 maternal haplotype centromeric regions and their lengths.**

**Table S9 CQ v3.2 paternal haplotype centromeric regions and their lengths.**

**Table S10 CQ v3.2 maternal haplotype centromeric higher-order repeat coordinates.**

**Table S11 CQ v3.2 paternal haplotype centromeric higher-order repeat coordinates.**

**Table S12 Higher-order repeat copies of the chromosome 1 centromeric region, compared among the CHM13, HG002, CN1, CQ, and YAO genomes.**

**Table S13 Contig N50 and complete centromere statistics in the LCL5/LCL6 genomes assembled with hifiasm and verkko at different ONT coverage levels ( > 100 kb).**

**Table S14 Complete centromere length distributions (bp) and maximum size variation in the LCL5/LCL6 genomes assembled with hifiasm and verkko at different ONT ( > 100 kb) coverage depths.**

## References

[1] Nurk S, Koren S, Rhie A, Rautiainen M, Bzikadze AV, Mikheenko A, et al. The complete sequence of a human genome. Science 2022;376(6588):44–53.

[2] Miga KH. Centromere studies in the era of ’telomere-to-telomere’ genomics. Exp Cell Res 2020;394(2):112127.

[3] Logsdon GA, Rozanski AN, Ryabov F, Potapova T, Shepelev VA, Catacchio CR, et al. The variation and evolution of complete human centromeres. Nature 2024;629(8010):136–45.

[4] Porubsky D, Dashnow H, Sasani TA, Logsdon GA, Hallast P, Noyes MD, et al. Human de novo mutation rates from a four-generation pedigree reference. Nature 2025;doi:10.1038/s41586-025-08922-2.

[5] Wang J, Wang W, Li R, Li Y, Tian G, Goodman L, et al. The diploid genome sequence of an Asian individual. Nature 2008;456(7218):60-5.

[6] Shi L, Guo Y, Dong C, Huddleston J, Yang H, Han X, et al. Long-read sequencing and de novo assembly of a Chinese genome. Nat Commun 2016;7:12065.

[7] Du Z, Ma L, Qu H, Chen W, Zhang B, Lu X, et al. Whole Genome Analyses of Chinese Population and De Novo Assembly of A Northern Han Genome. Genomics Proteomics Bioinformatics 2019;17(3):229–47.

[8] Chao KH, Zimin AV, Pertea M, Salzberg SL. The first gapless, reference-quality, fully annotated genome from a Southern Han Chinese individual. G3 (Bethesda) 2023;13(3).

[9] Yang X, Zhao X, Qu S, Jia P, Wang B, Gao S, et al. Haplotype-resolved Chinese male genome assembly based on high-fidelity sequencing. Fundam Res 2022;2(6):946–53.

[10] Yang C, Zhou Y, Song Y, Wu D, Zeng Y, Nie L, et al. The complete and fully-phased diploid genome of a male Han Chinese. Cell Res 2023;33(10):745–61.

[11] He Y, Chu Y, Guo S, Hu J, Li R, Zheng Y, et al. T2T-YAO: A Telomere-to-telomere Assembled Diploid Reference Genome for Han Chinese. Genomics Proteomics Bioinformatics 2023;21(6):1085–100.

[12] Logsdon GA, Ebert P, Audano PA, Loftus M, Porubsky D, Ebler J, et al. Complex genetic variation in nearly complete human genomes. bioRxiv 2024;614721.

[13] Zheng Y, Liu Y, Yang J, Dong L, Zhang R, Tian S, et al. Multi-omics data integration using ratio-based quantitative profiling with Quartet reference materials. Nat Biotechnol 2024;42(7):1133–49.

[14] Ren L, Duan X, Dong L, Zhang R, Yang J, Gao Y, et al. Quartet DNA reference materials and datasets for comprehensively evaluating germline variant calling performance. Genome Biol 2023;24(1):270.

[15] Yang J, Liu Y, Shang J, Chen Q, Chen Q, Ren L, et al. The Quartet Data Portal: integration of community-wide resources for multiomics quality control. Genome Biol 2023;24(1):245.

[16] Jia P, Dong L, Yang X, Wang B, Bush SJ, Wang T, et al. Haplotype-resolved assemblies and variant benchmark of a Chinese Quartet. Genome Biol 2023;24(1):277.

[17] Ren L, Shi L, Zheng Y. Reference Materials for Improving Reliability of Multiomics Profiling. Phenomics 2024;4(5):487–521.

[18] Zhang N, Chen Q, Zhang P, Zhou K, Liu Y, Wang H, et al. Quartet metabolite reference materials for inter-laboratory proficiency test and data integration of metabolomics profiling. Genome Biol 2024;25(1):34.

[19] Talbert PB, Henikoff S. What makes a centromere? Exp Cell Res 2020;389(2):111895.

[20] Logsdon GA, Vollger MR, Hsieh P, Mao Y, Liskovykh MA, Koren S, et al. The structure, function and evolution of a complete human chromosome 8. Nature 2021;593(7857):101-7.

[21] McKinley KL, Cheeseman IM. The molecular basis for centromere identity and function. Nat Rev Mol Cell Biol 2016;17(1):16–29.

[22] Rautiainen M, Nurk S, Walenz BP, Logsdon GA, Porubsky D, Rhie A, et al. Telomere-to-telomere assembly of diploid chromosomes with Verkko. Nat Biotechnol 2023;41(10):1474–82.

[23] Antipov D, Rautiainen M, Nurk S, Walenz BP, Solar SJ, Phillippy AM, et al. Verkko2 integrates proximity-ligation data with long-read De Bruijn graphs for efficient telomere-to-telomere genome assembly, phasing, and scaffolding. Genome Res 2025;35(7):1583–94.

[24] Cheng H, Asri M, Lucas J, Koren S, Li H. Scalable telomere-to-telomere assembly for diploid and polyploid genomes with double graph. Nat Methods 2024;21(6):967–70.

[25] Xu M, Guo L, Gu S, Wang O, Zhang R, Peters BA, et al. TGS-GapCloser: a fast and accurate gap closer for large genomes with low coverage of error-prone long reads. GigaScience 2020;9(9):giaa094.

[26] Rautiainen M. Ribotin: automated assembly and phasing of rDNA morphs. Bioinformatics 2024;40(3):btae124.

[27] Hu J, Wang Z, Liang F, Liu SL, Ye K, Wang DP. NextPolish2: a repeat-aware polishing tool for genomes assembled using HiFi long reads. Genomics Proteomics Bioinformatics 2024;22(1):qzad009.

[28] Mc Cartney AM, Shafin K, Alonge M, Bzikadze AV, Formenti G, Fungtammasan A, et al. Chasing perfection: validation and polishing strategies for telomere-to-telomere genome assemblies. Nat Methods 2022;19(6):687–95.

[29] Rhie A, Walenz BP, Koren S, Phillippy AM. Merqury: reference-free quality, completeness, and phasing assessment for genome assemblies. Genome Biol 2020;21(1):245.

[30] Patterson M, Marschall T, Pisanti N, Van Iersel L, Stougie L, Klau GW, et al. WhatsHap: weighted haplotype assembly for future-generation sequencing reads. Journal of Computational Biology 2015;22(6):498–509.

[31] Cheng H, Concepcion GT, Feng X, Zhang H, Li H. Haplotype-resolved de novo assembly using phased assembly graphs with hifiasm. Nat Methods 2021;18(2):170–5.

[32] Liao WW, Asri M, Ebler J, Doerr D, Haukness M, Hickey G, et al. A draft human pangenome reference. Nature 2023;617(7960):312-24.

[33] Chen Q, Yang C, Zhang G, Wu D. GCI: a continuity inspector for complete genome assembly. Bioinformatics 2024;40(11):btae633.

[34] Li K, Xu P, Wang J, Yi X, Jiao Y. Identification of errors in draft genome assemblies at single-nucleotide resolution for quality assessment and improvement. Nat Commun 2023;14(1):6556.

[35] Goel M, Sun H, Jiao W-B, Schneeberger K. SyRI: finding genomic rearrangements and local sequence differences from whole-genome assemblies. Genome Biol 2019;20:1–13.

[36] Vollger MR, Kerpedjiev P, Phillippy AM, Eichler EE. StainedGlass: interactive visualization of massive tandem repeat structures with identity heatmaps. Bioinformatics 2022;38(7):2049–51.

[37] Gao S, Zhang Y, Bush SJ, Wang B, Yang X, Ye K. Centromere landscapes resolved from hundreds of human genomes. Genomics Proteomics Bioinformatics 2024;qzae071.

[38] Taylor DJ, Eizenga JM, Li Q, Das A, Jenike KM, Kenny EE, et al. Beyond the Human Genome Project: The age of complete human genome sequences and pangenome references. Annu Rev Genoma Hum G 2024;25.

[39] Wang B, Dang N, Yang X, Xu S, Ye K. The human pangenome reference: the beginning of a new era for genomics. Sci Bull (Beijing) 2023;68(14):1484–7.

[40] Pan B, Ren L, Onuchic V, Guan M, Kusko R, Bruinsma S, et al. Assessing reproducibility of inherited variants detected with short-read whole genome sequencing. Genome Biol 2022;23(1):2.

[41] Wang B, Yang X, Jia Y, Xu Y, Jia P, Dang N, et al. High-quality Arabidopsis thaliana Genome Assembly with Nanopore and HiFi Long Reads. Genomics Proteomics Bioinformatics 2022;20(1):4–13.

[42] Gong G, Dan C, Xiao S, Guo W, Huang P, Xiong Y, et al. Chromosomal-level assembly of yellow catfish genome using third-generation DNA sequencing and Hi-C analysis. GigaScience 2018;7(11):giy120.

[43] Durand NC, Shamim MS, Machol I, Rao SS, Huntley MH, Lander ES, et al. Juicer provides a one-click system for analyzing loop-resolution Hi-C experiments. Cell Syst 2016;3(1):95–8.

[44] Durand NC, Robinson JT, Shamim MS, Machol I, Mesirov JP, Lander ES, et al. Juicebox provides a visualization system for Hi-C contact maps with unlimited zoom. Cell Syst 2016;3(1):99–101.

[45] van Dongen J, Slagboom PE, Draisma HH, Martin NG, Boomsma DI. The continuing value of twin studies in the omics era. Nat Rev Genet 2012;13(9):640–53.

[46] Nurk S, Walenz BP, Rhie A, Vollger MR, Logsdon GA, Grothe R, et al. HiCanu: accurate assembly of segmental duplications, satellites, and allelic variants from high-fidelity long reads. Genome Res 2020;30(9):1291–305.

[47] Wick RR, Judd LM, Gorrie CL, Holt KE. Unicycler: Resolving bacterial genome assemblies from short and long sequencing reads. PLoS Comput Biol 2017;13(6):e1005595.

[48] Gao Y, Yang X, Chen H, Tan X, Yang Z, Deng L, et al. A pangenome reference of 36 Chinese populations. Nature 2023;619(7968):112-21.

[49] Li H. Minimap2: pairwise alignment for nucleotide sequences. Bioinformatics 2018;34(18):3094–100.

[50] Marçais G, Delcher AL, Phillippy AM, Coston R, Salzberg SL, Zimin A. MUMmer4: A fast and versatile genome alignment system. PLoS Comput Biol 2018;14(1):e1005944.

[51] Zhou Z, Yu Z, Huang X, Liu J, Guo Y, Chen L, et al. GenomeSyn: a bioinformatics tool for visualizing genome synteny and structural variations. J Genet Genomics 2022;49(12):1174–6.

[52] Koboldt DC, Zhang Q, Larson DE, Shen D, McLellan MD, Lin L, et al. VarScan 2: somatic mutation and copy number alteration discovery in cancer by exome sequencing. Genome Res 2012;22(3):568–76.

[53] Yoo D, Rhie A, Hebbar P, Antonacci F, Logsdon GA, Solar SJ, et al. Complete sequencing of ape genomes. Nature 2025;641(8062):401-18.

[54] Katoh K, Standley DM. MAFFT multiple sequence alignment software version 7: improvements in performance and usability. Mol Biol Evol 2013;30(4):772–80.

[55] Nguyen L-T, Schmidt HA, Von Haeseler A, Minh BQ. IQ-TREE: a fast and effective stochastic algorithm for estimating maximum-likelihood phylogenies. Mol Biol Evol 2015;32(1):268–74.

[56] Sanderson MJ. r8s: inferring absolute rates of molecular evolution and divergence times in the absence of a molecular clock. Bioinformatics 2003;19(2):301–2.

[57] Hall MB. Rasusa: Randomly subsample sequencing reads to a specified coverage. Journal of Open Source Software 2022;7(69):3941.

[58] Chen T, Chen X, Zhang S, Zhu J, Tang B, Wang A, et al. The genome sequence archive family: toward explosive data growth and diverse data types. Genomics Proteomics Bioinformatics 2021;19(4):578–83.

[59] Chen M, Ma Y, Wu S, Zheng X, Kang H, Sang J, et al. Genome Warehouse: A Public Repository Housing Genome-scale Data. Genomics Proteomics Bioinformatics 2021;19(4):584–9.

