## Supplementary figures and images for "A Telomere-to-Telomere Diploid Reference Genome and Centromere Structure of the Chinese Quartet"

### Figure S1

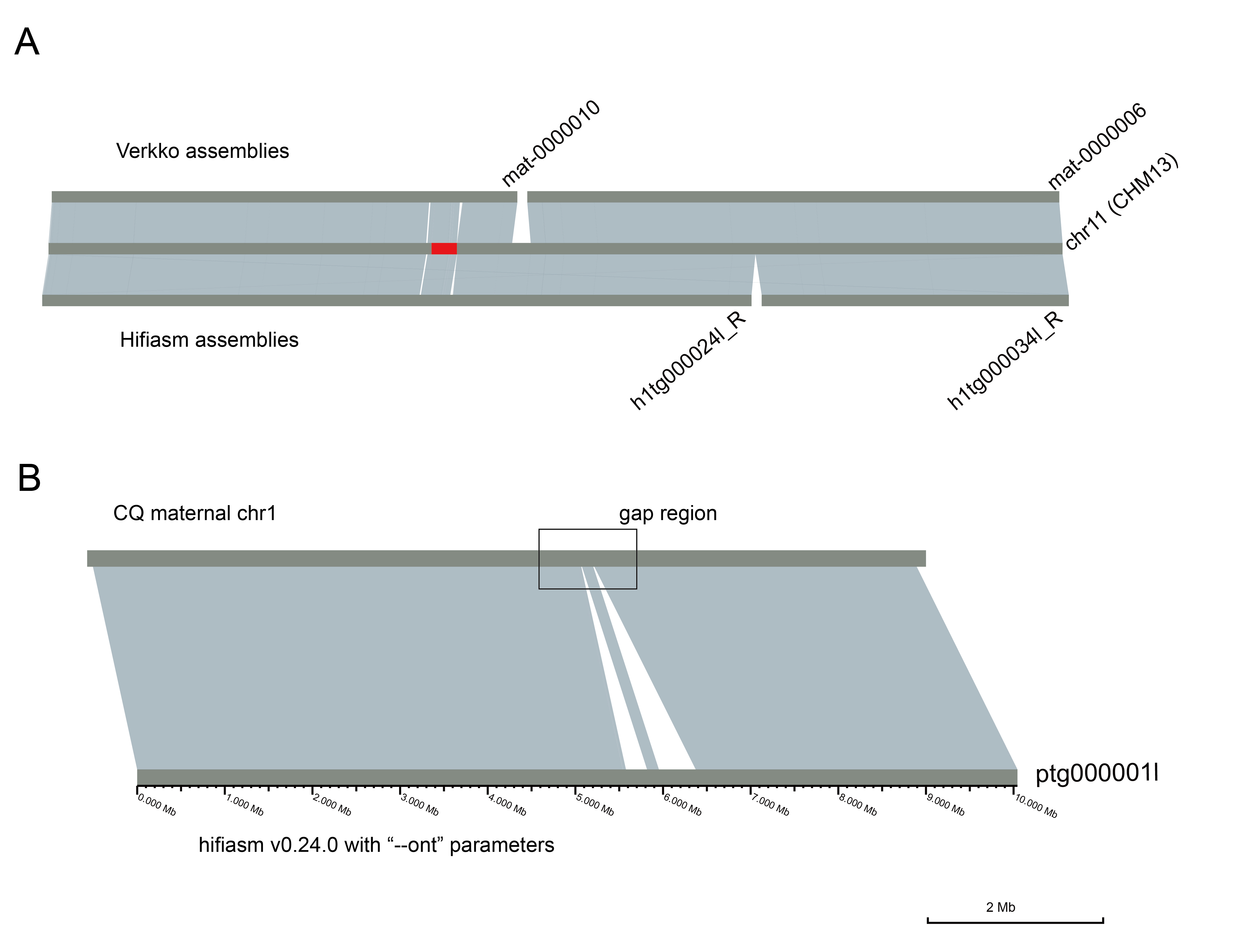

### Figure S2

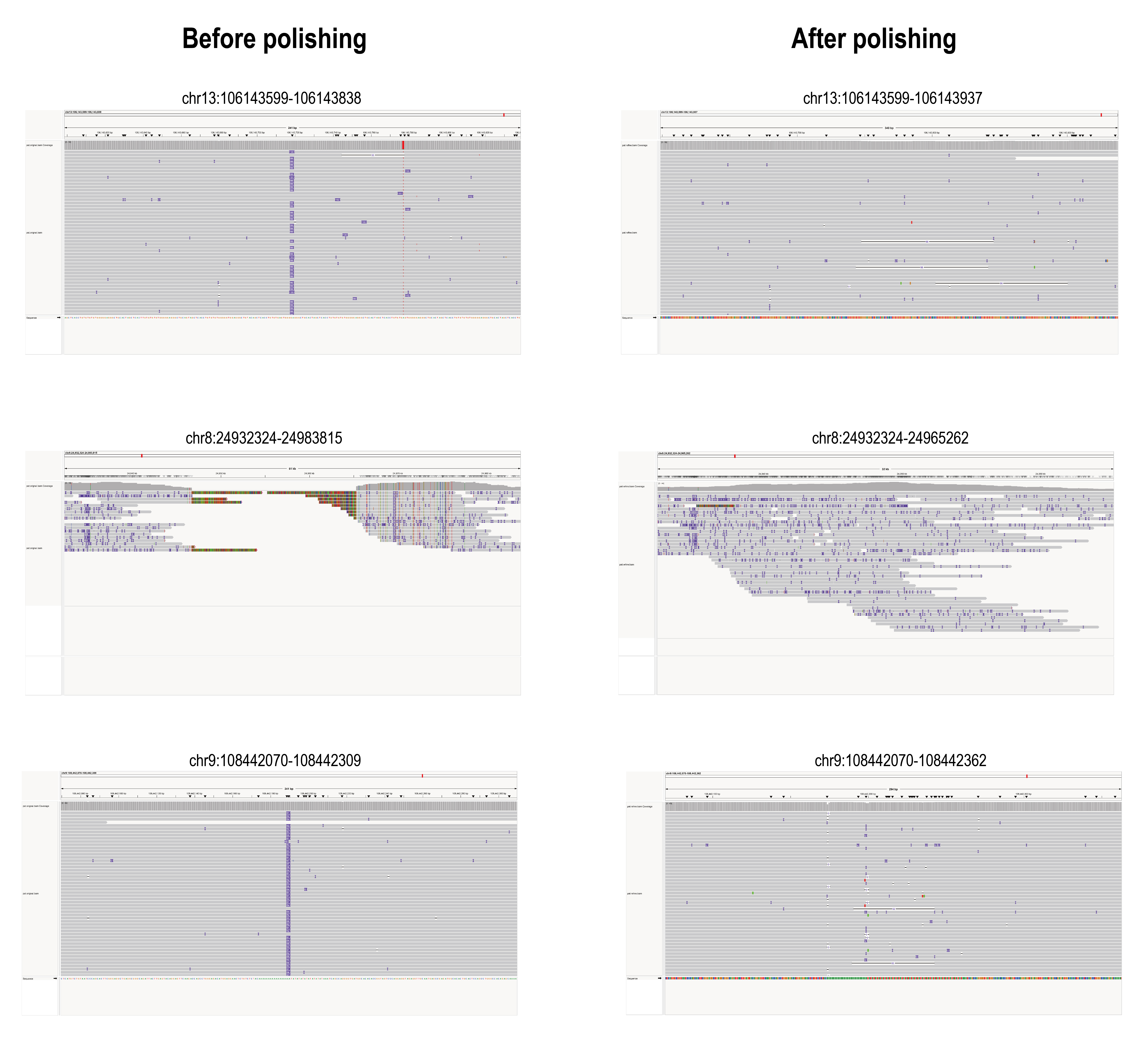

### Figure S3

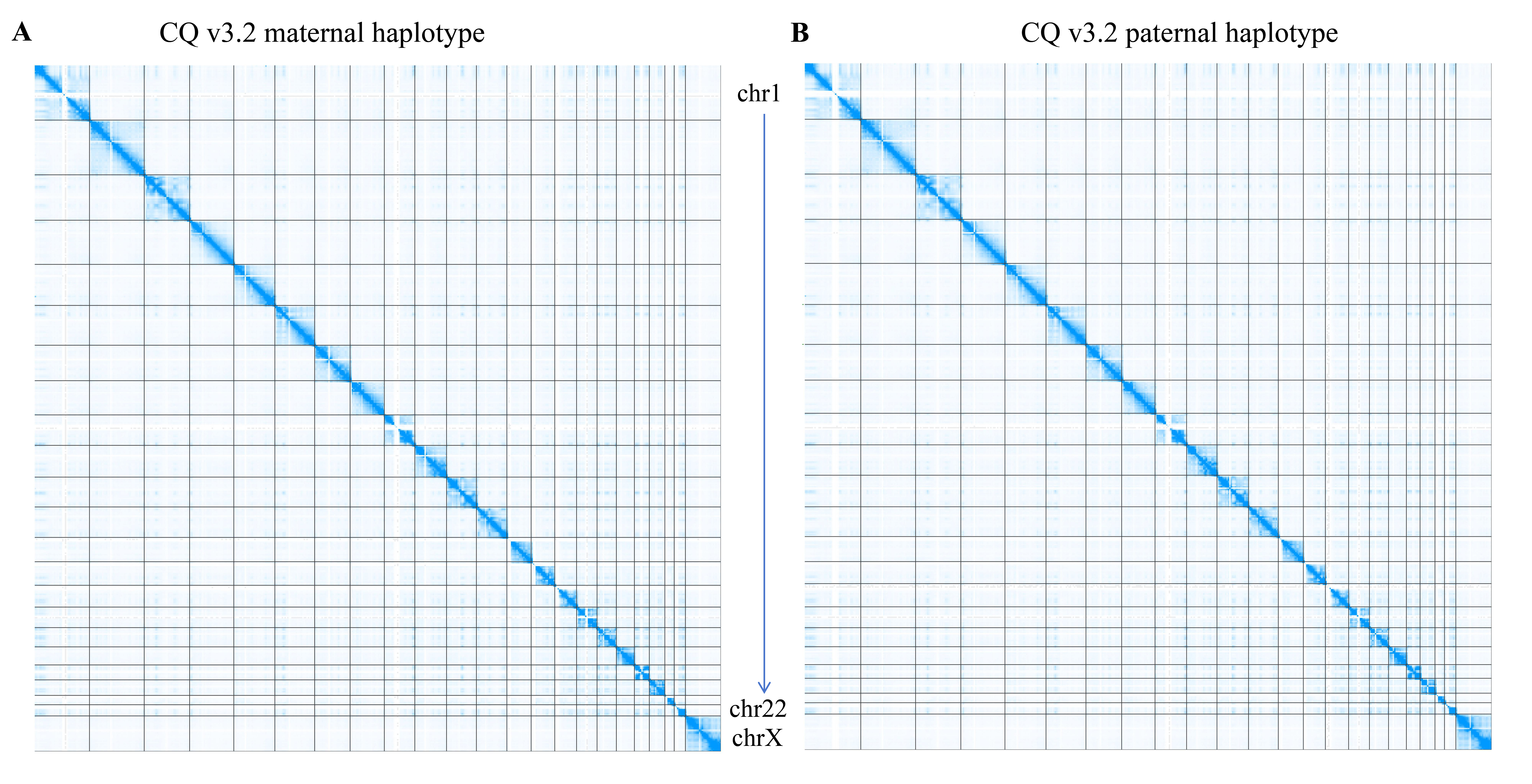

### Figure S4

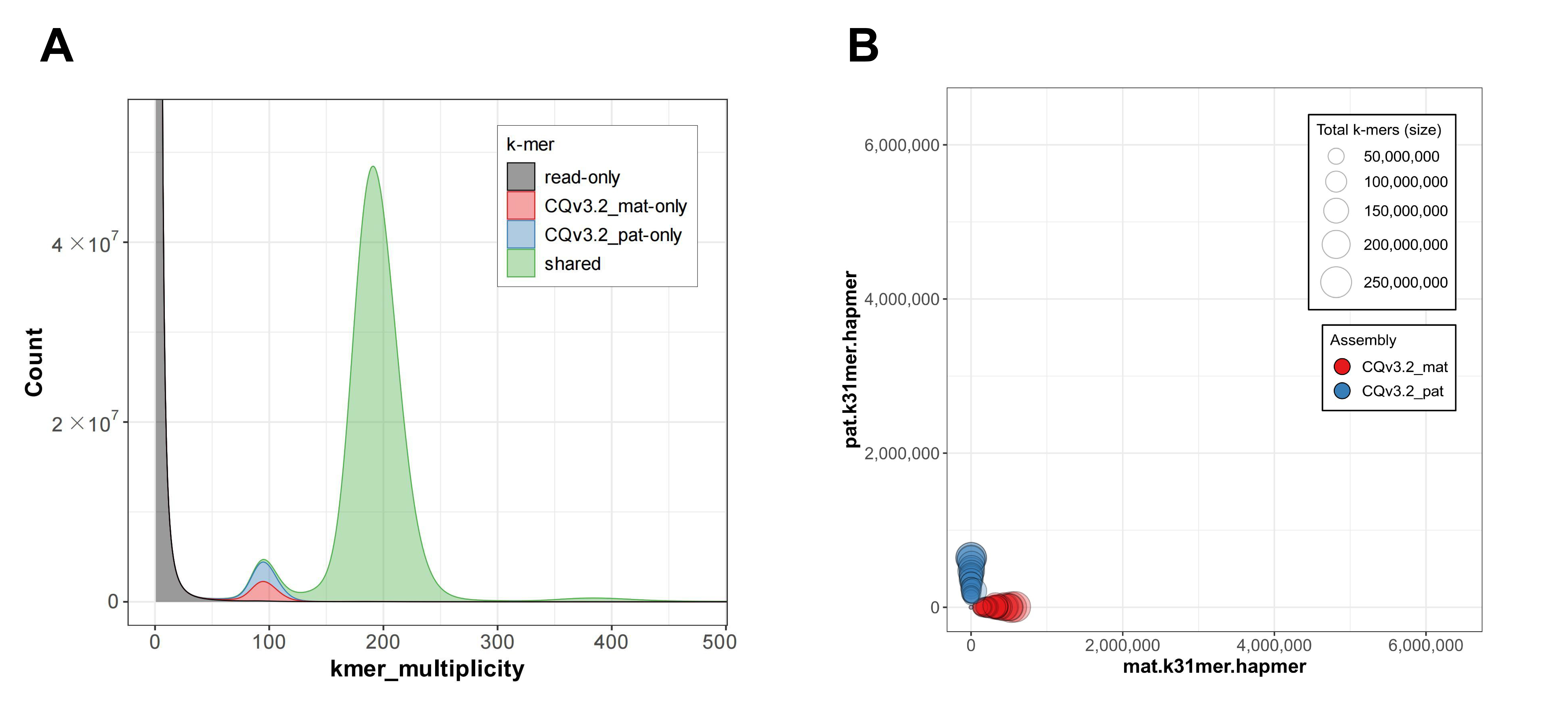

### Figure S5

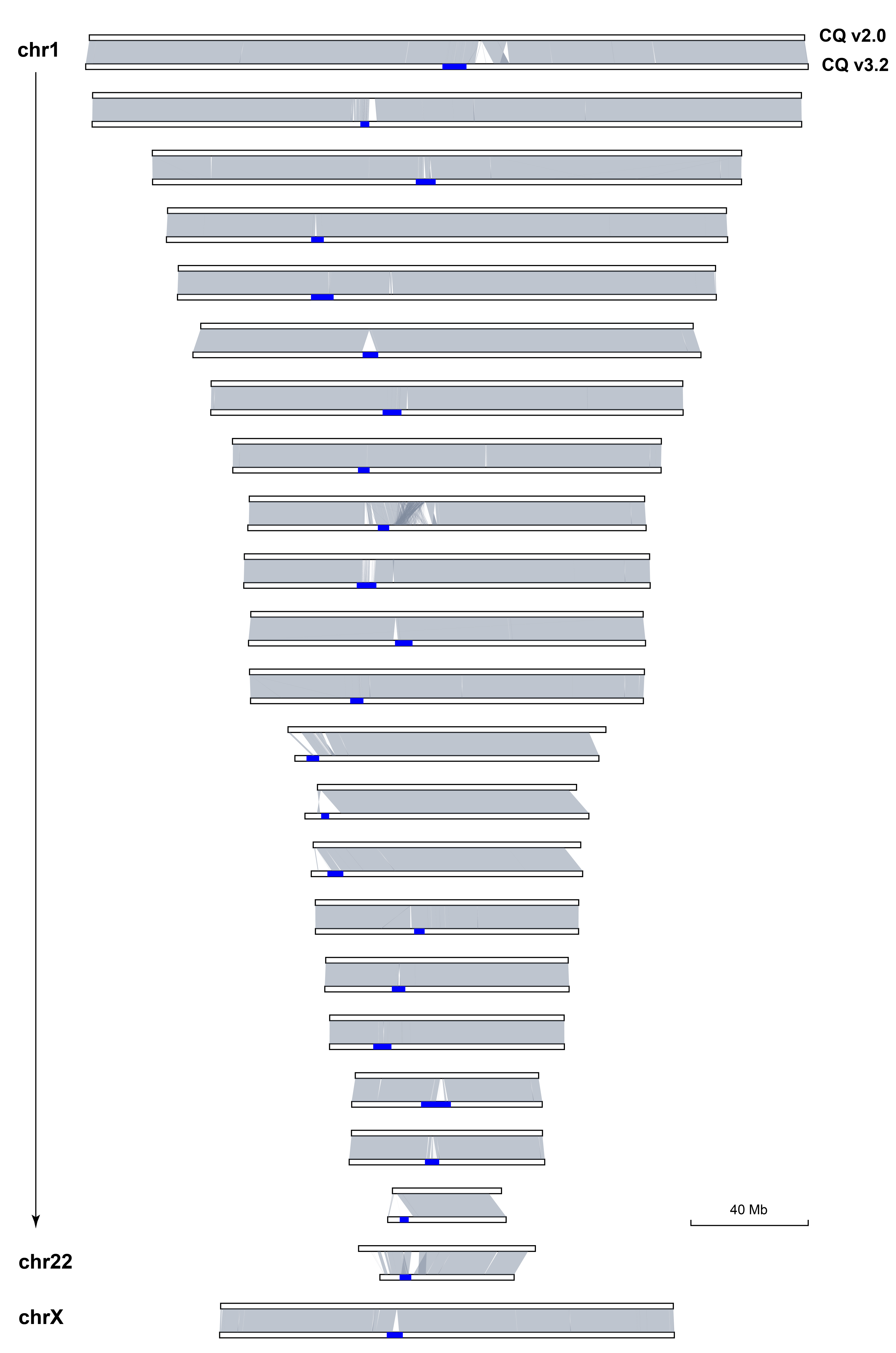

### Figure S6

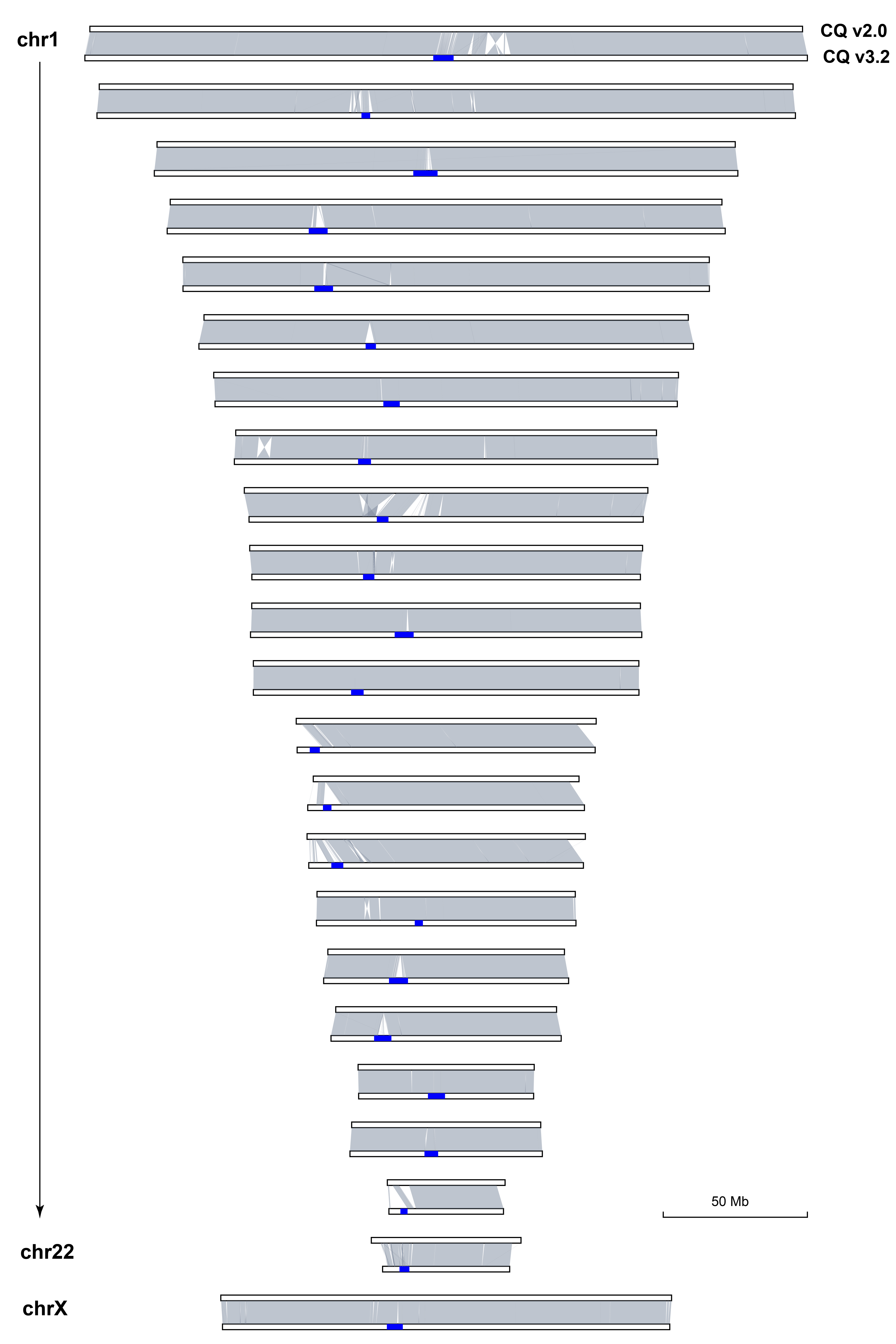

### Figure S7

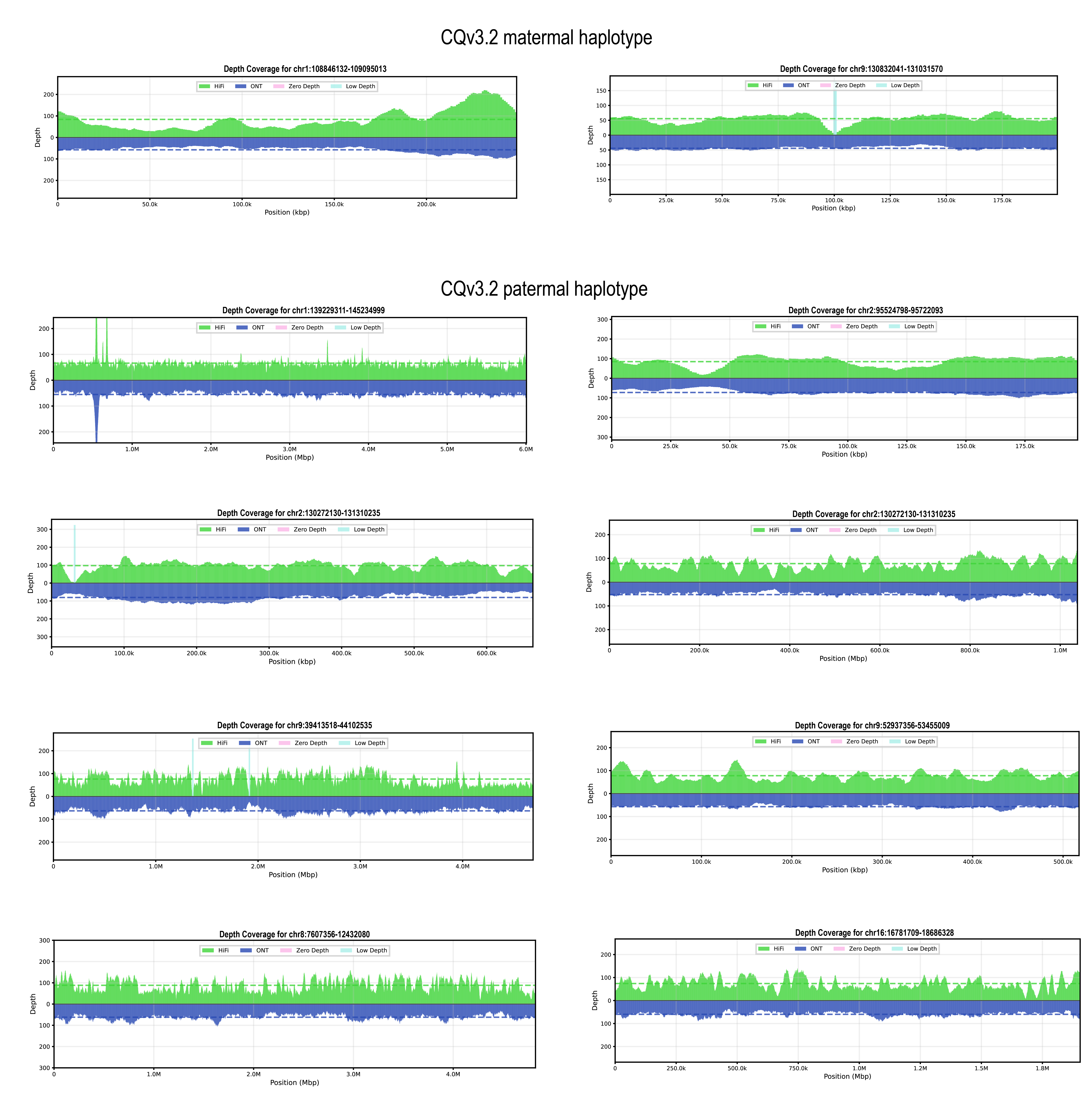

### Figure S8

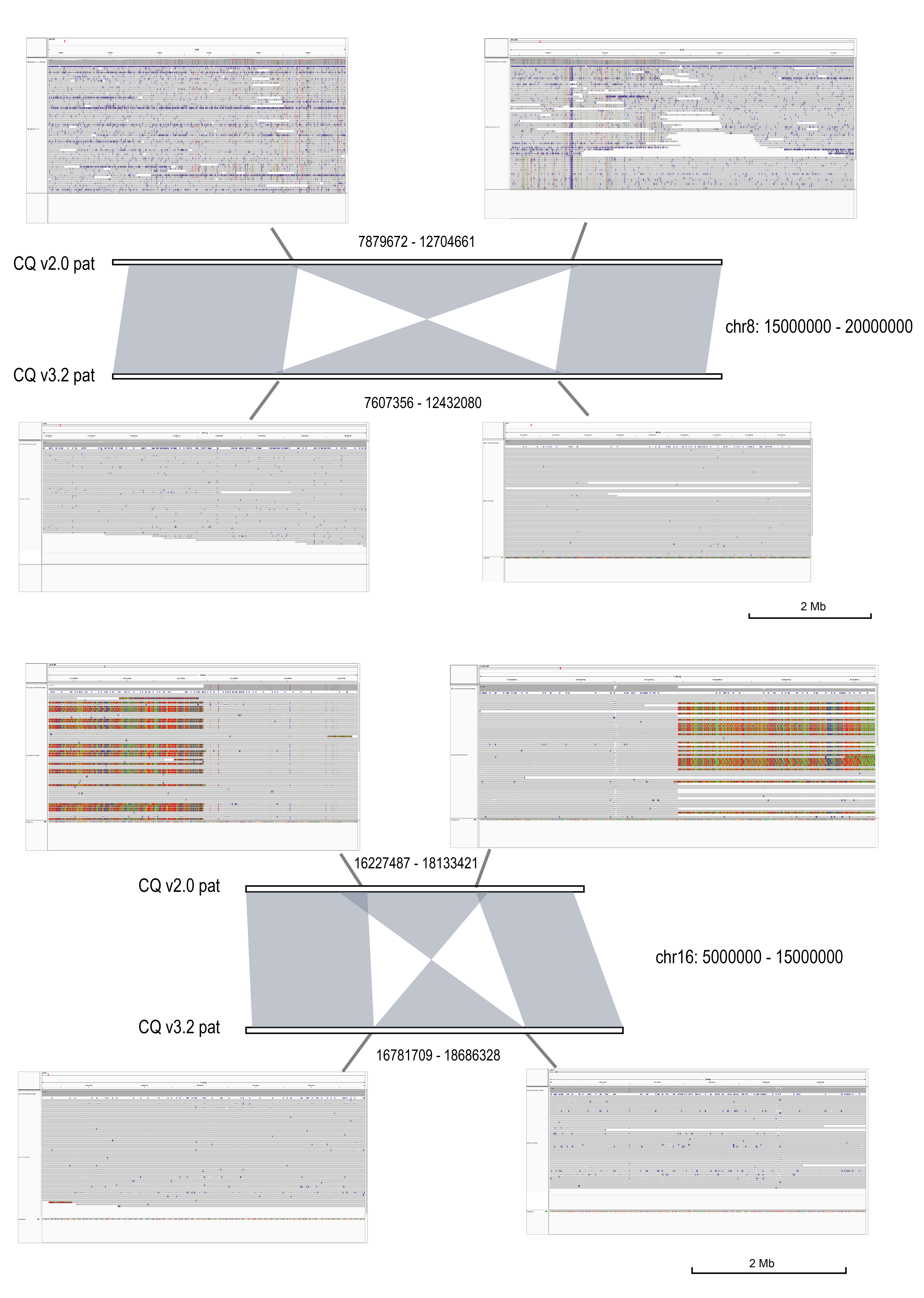

### Figure S9

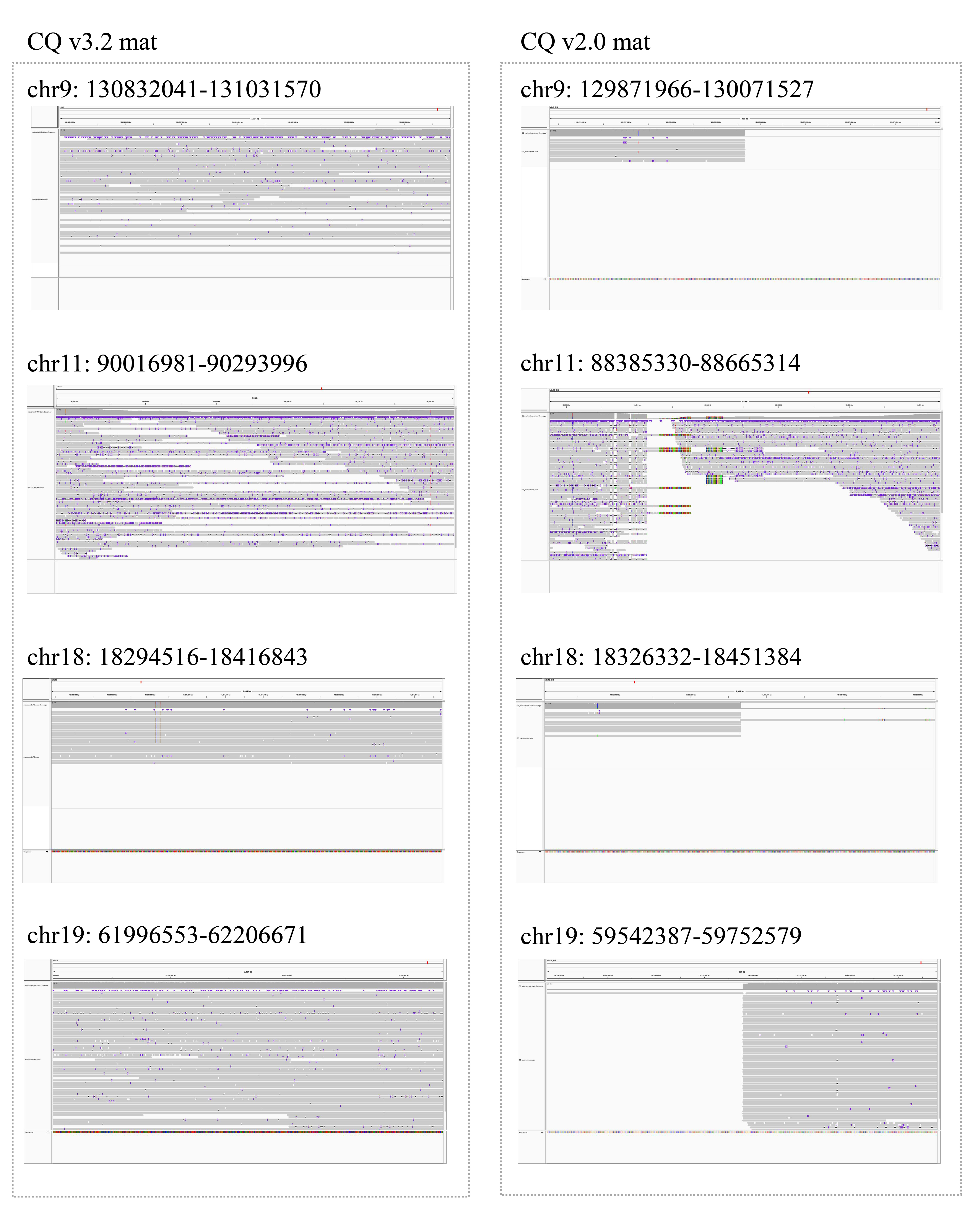

### Figure S10

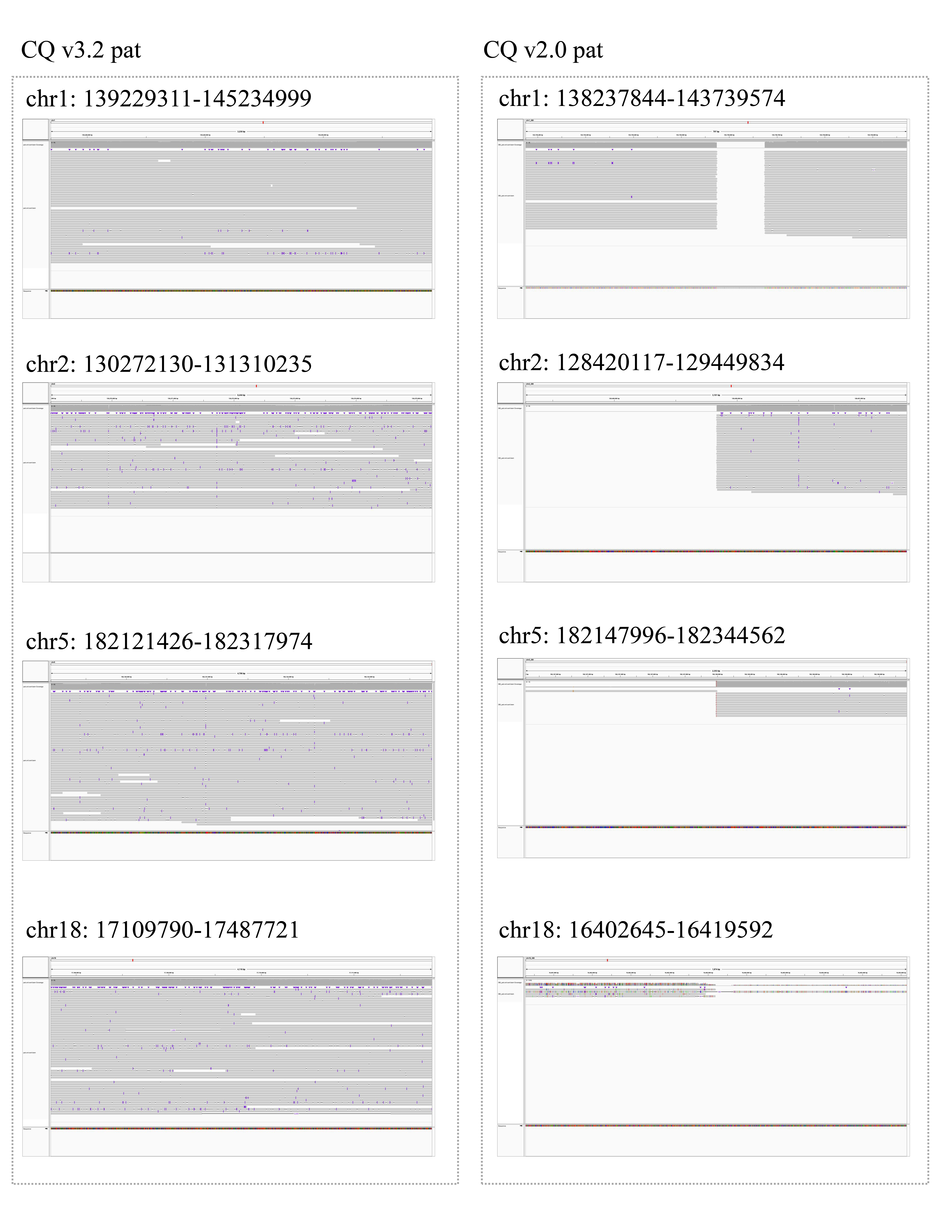

### Figure S11

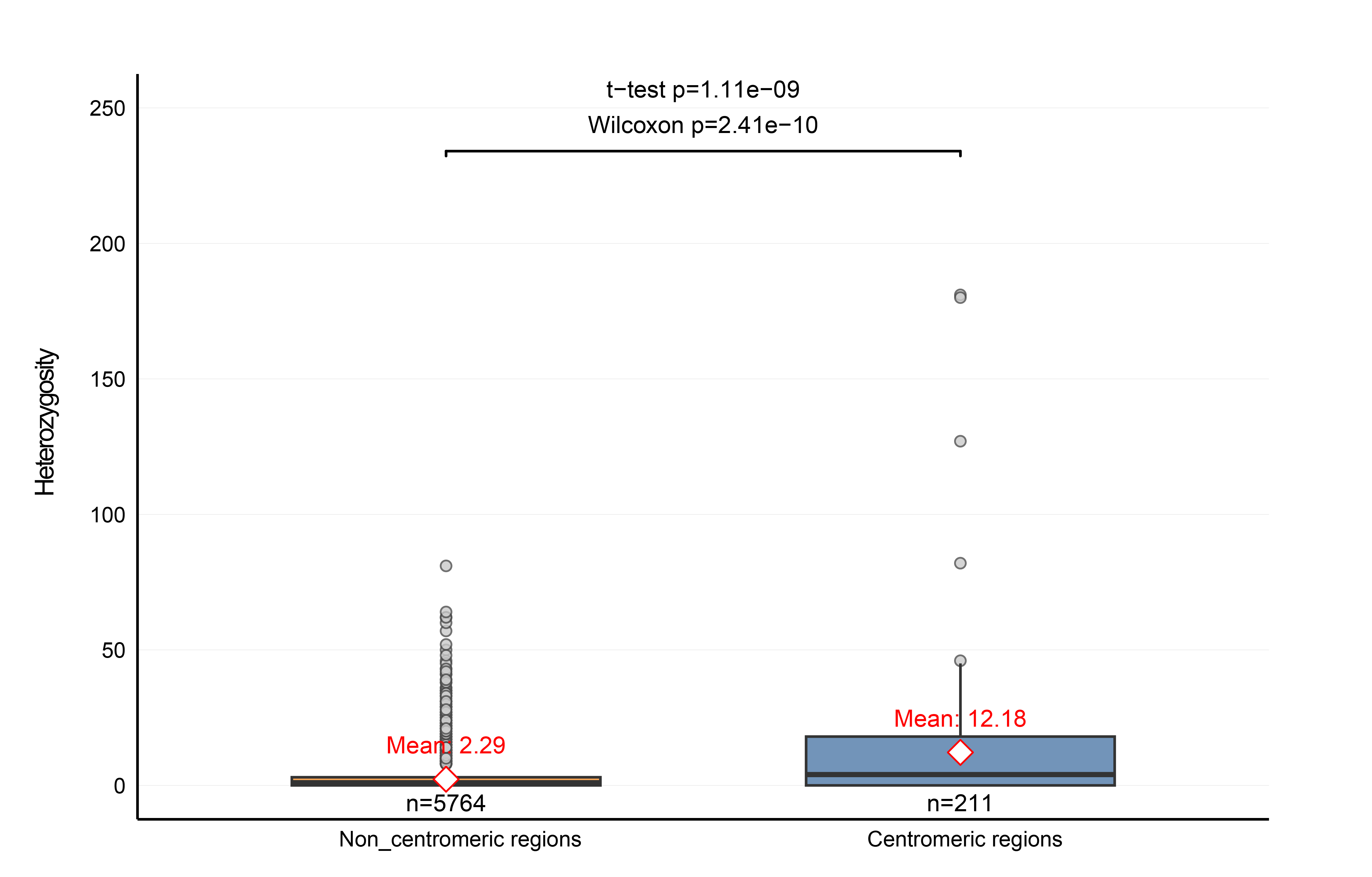

### Figure S12

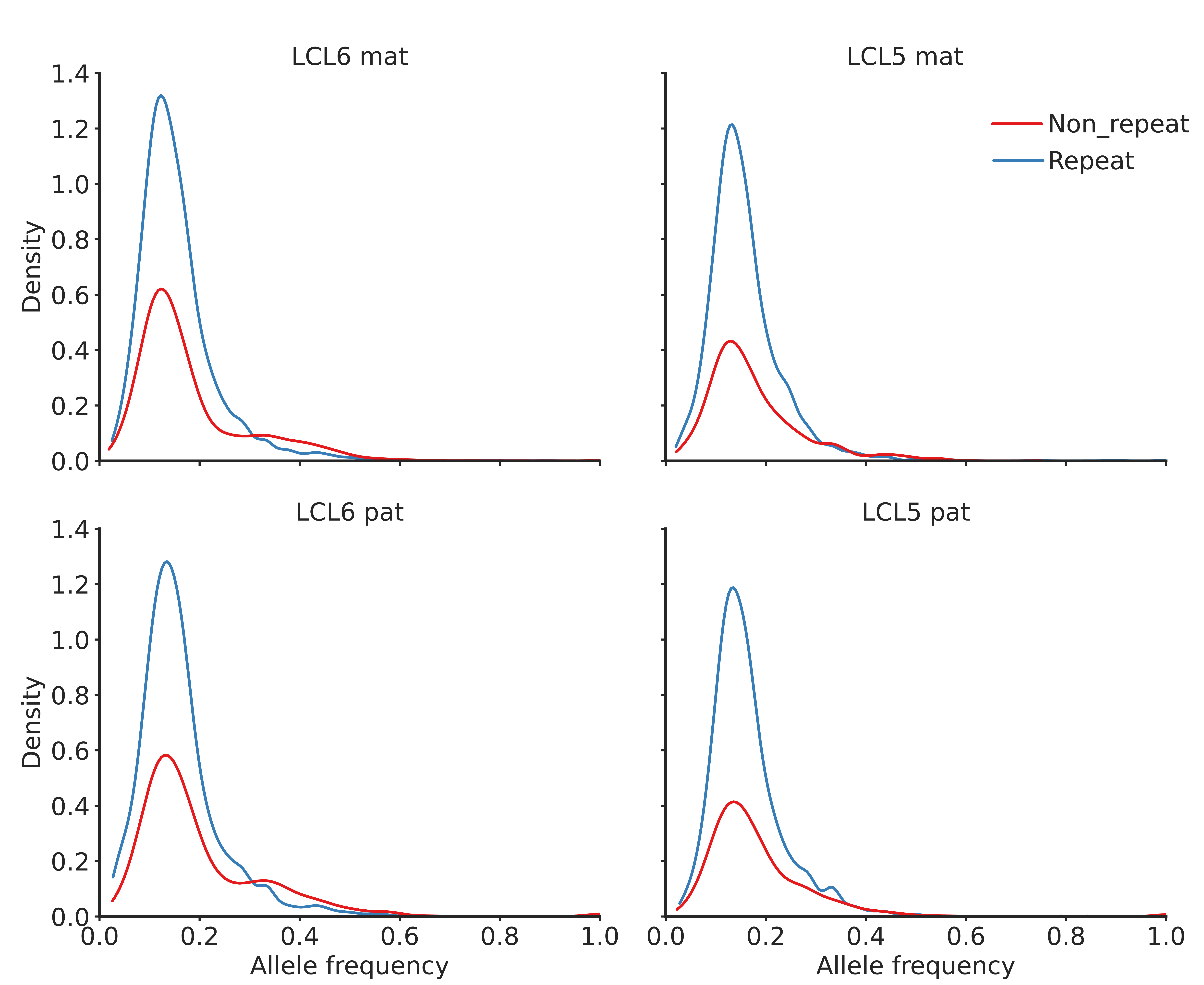

### Figure S13

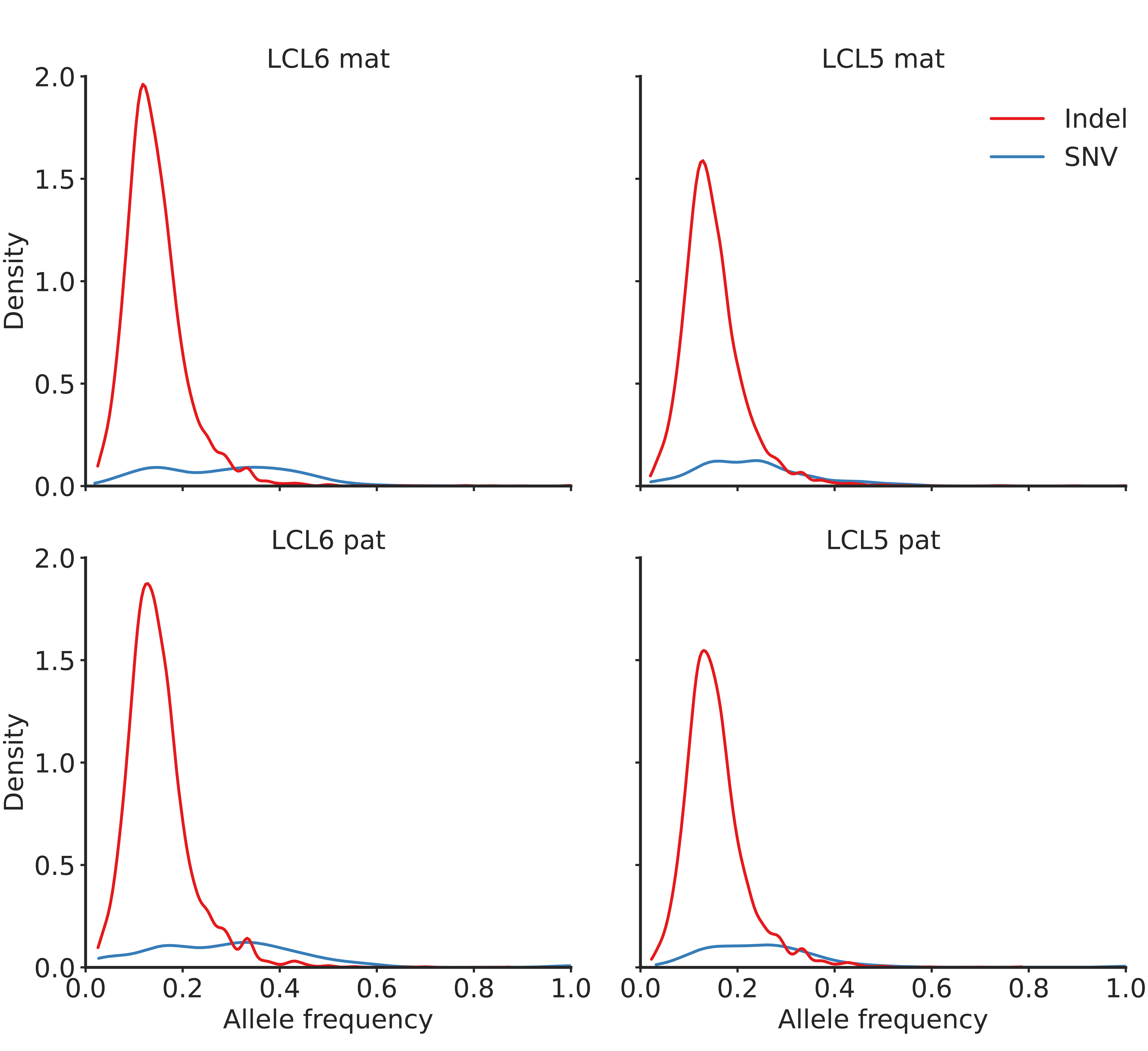

### Figure S14

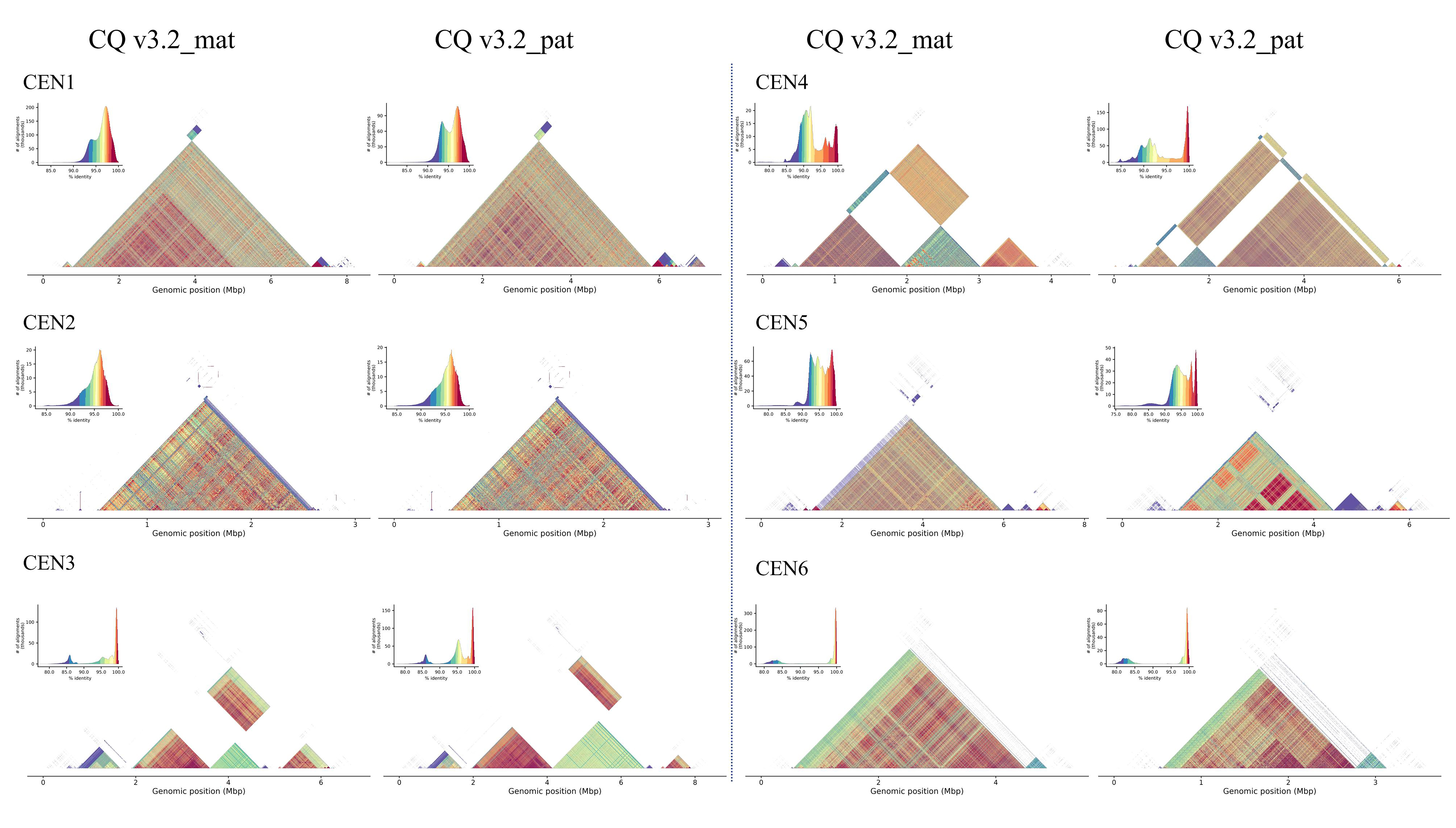

### Figure S15

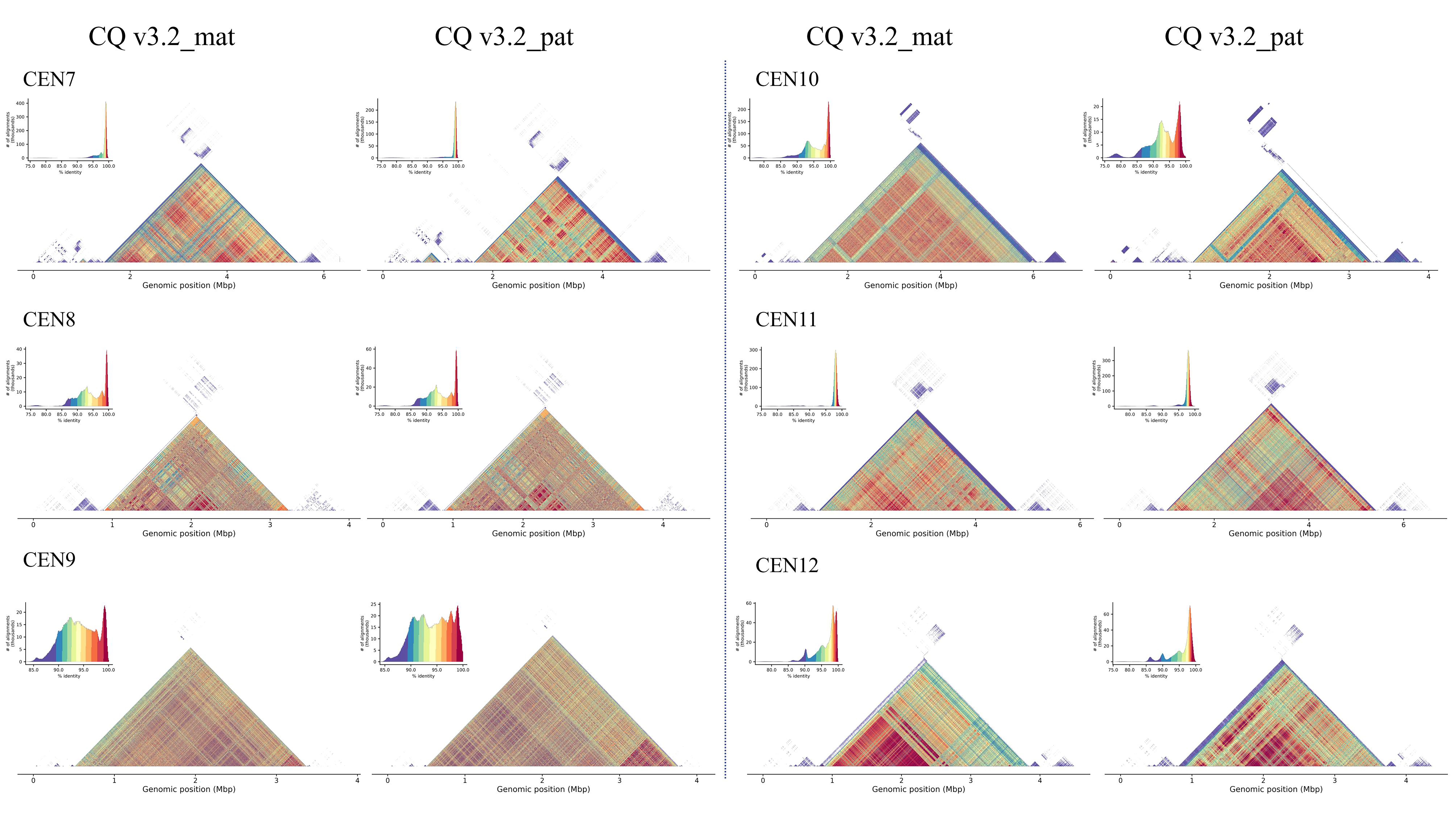

### Figure S16

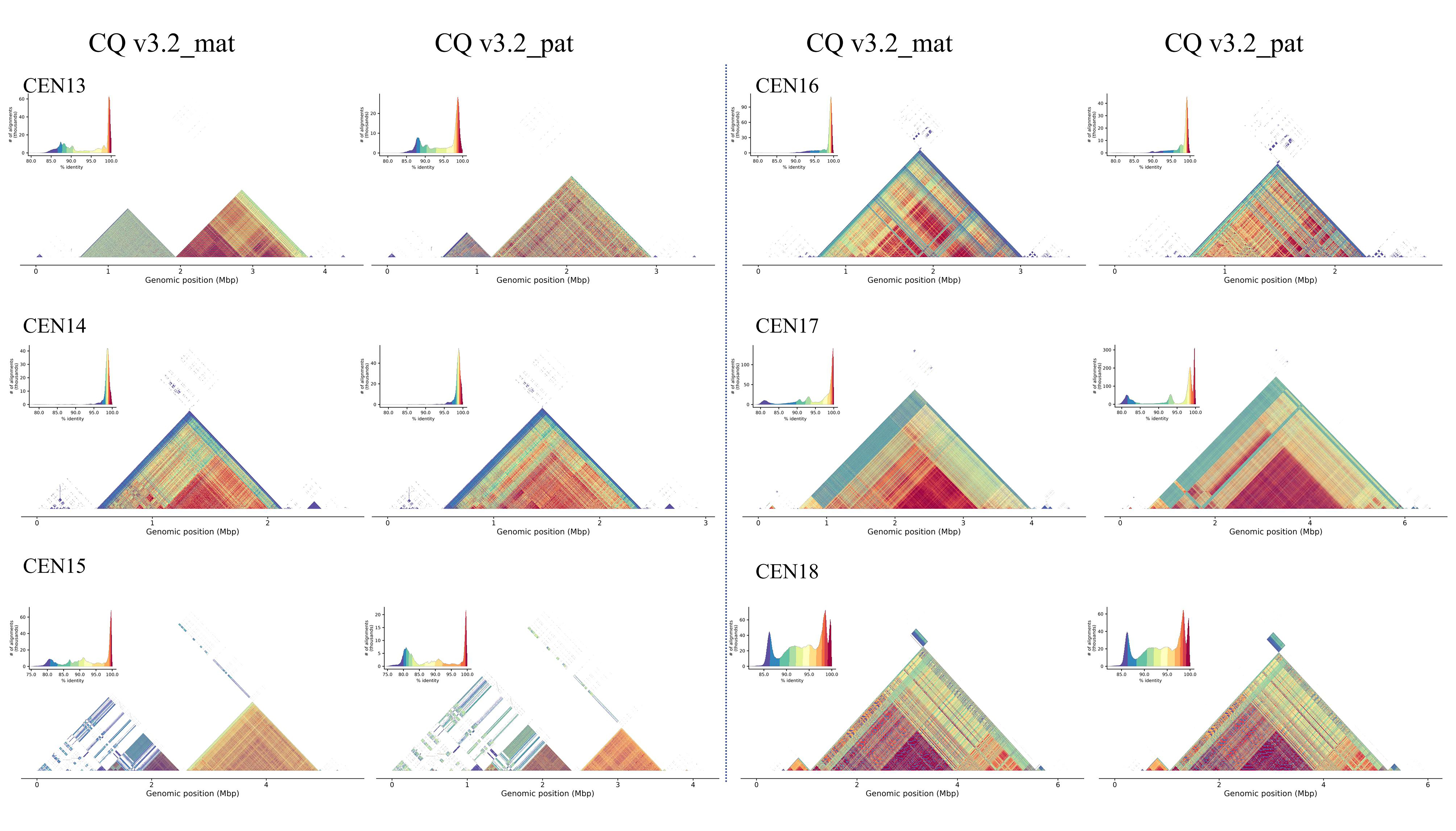

### Figure S17

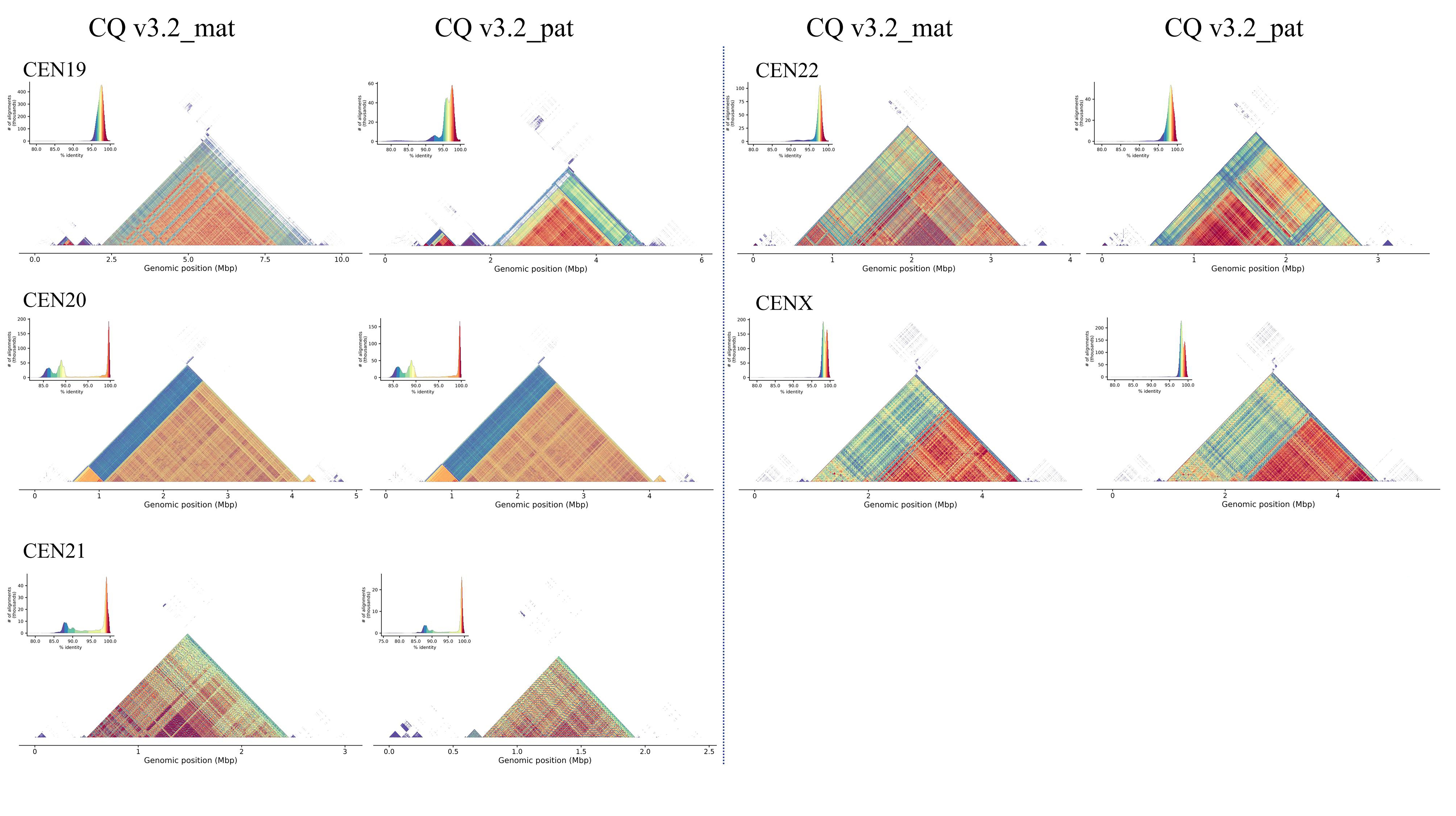

### Figure S18

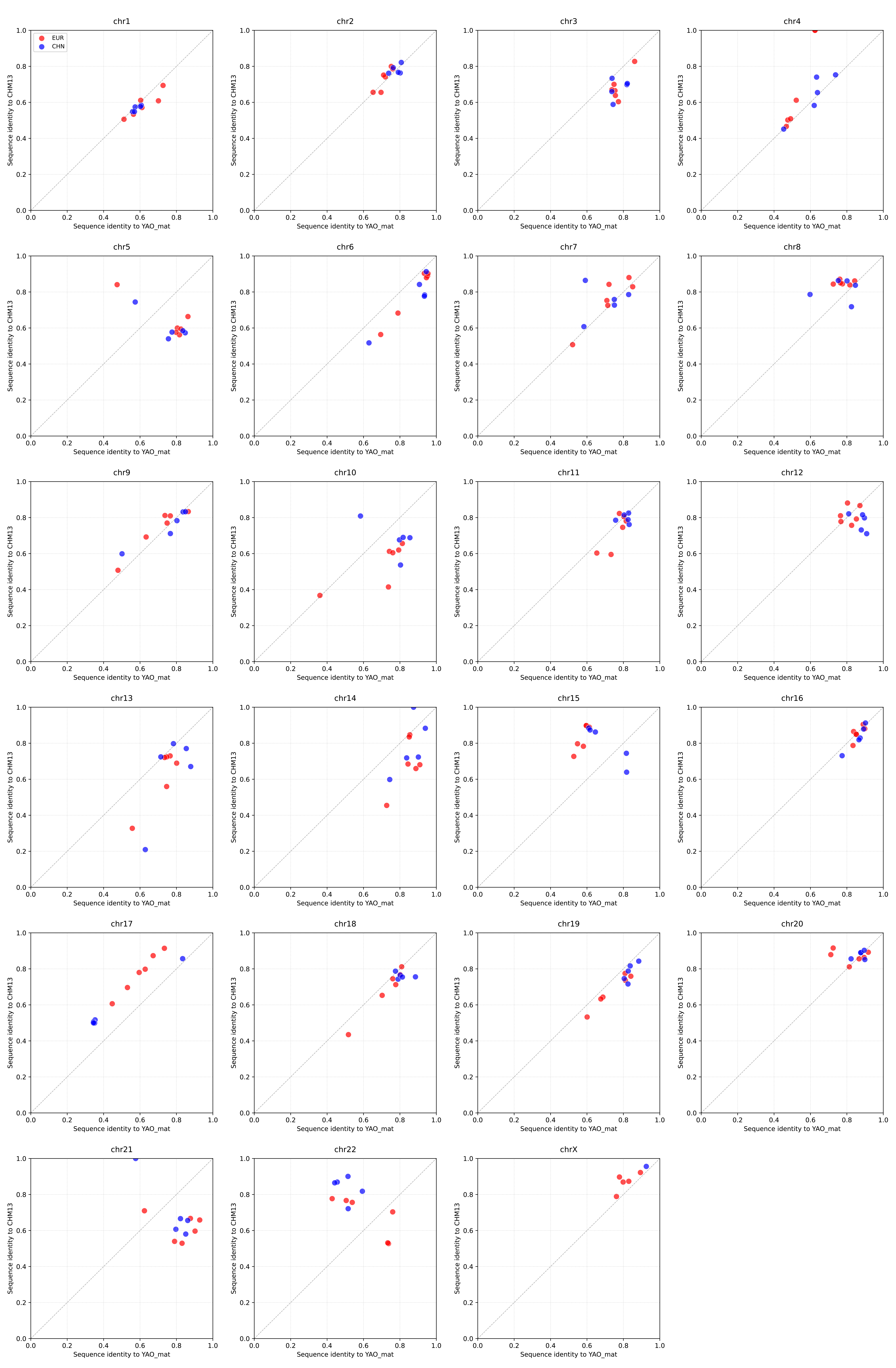

### Figure S19

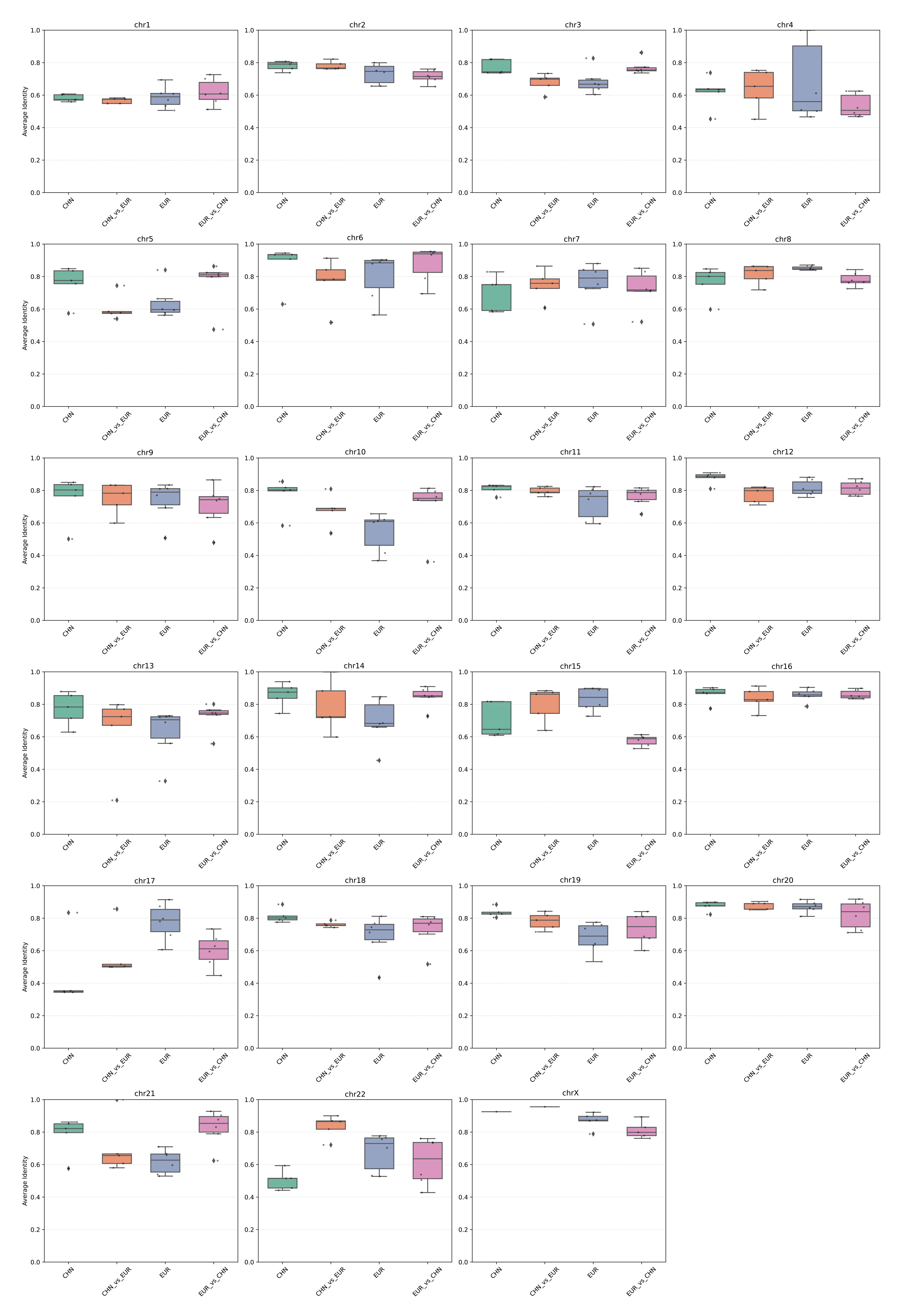

### Figure S20

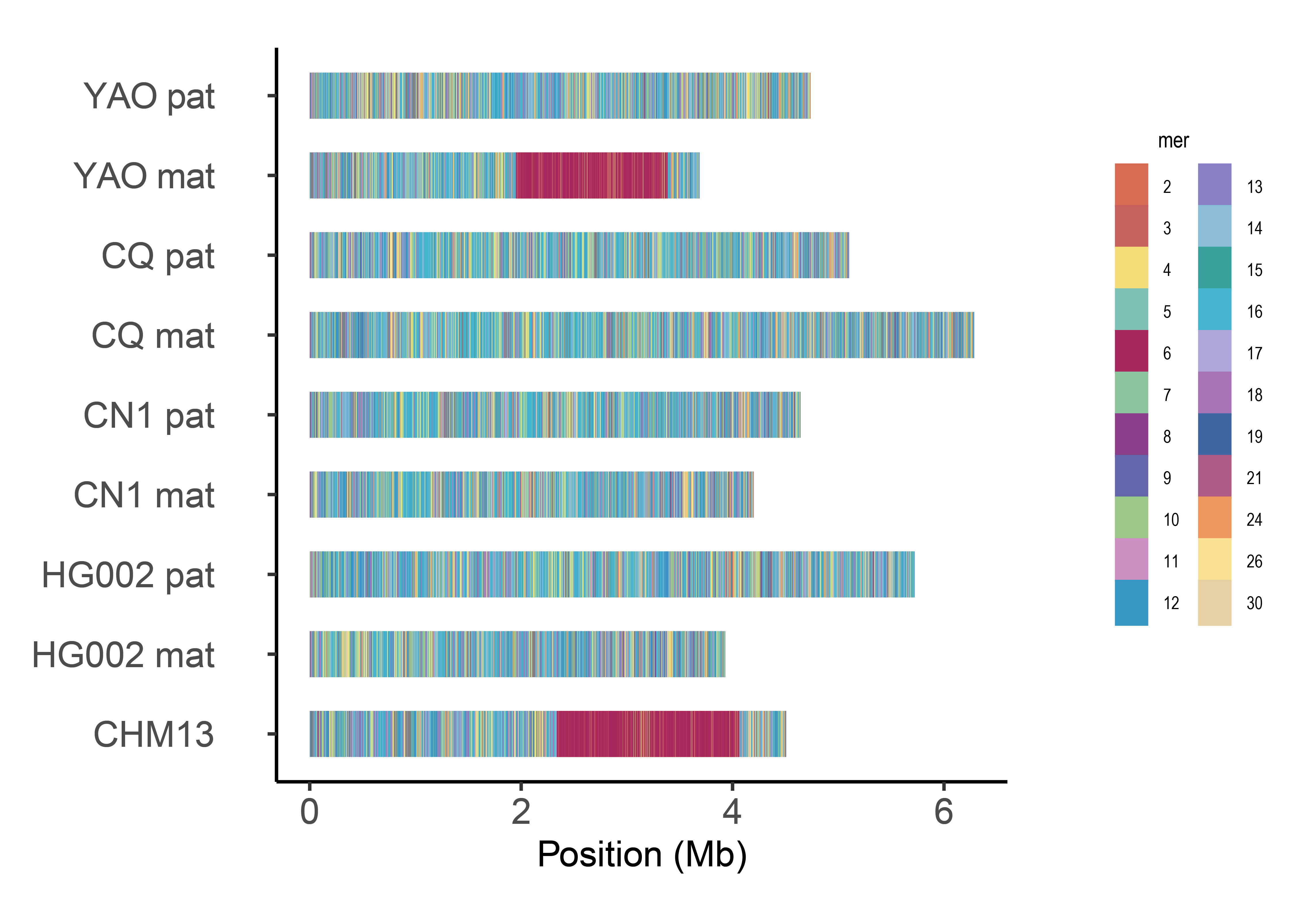

### Figure S21

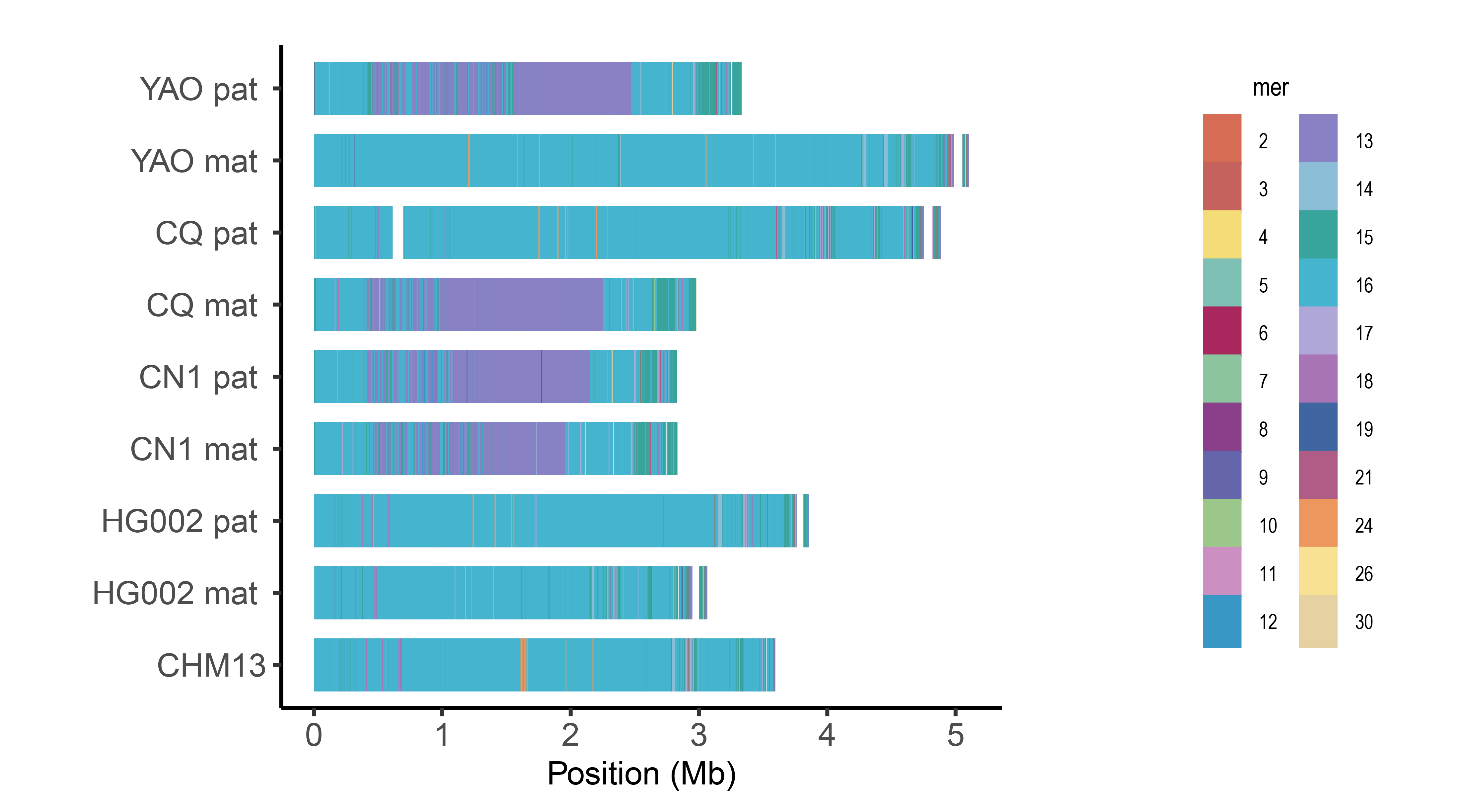

### Figure S22

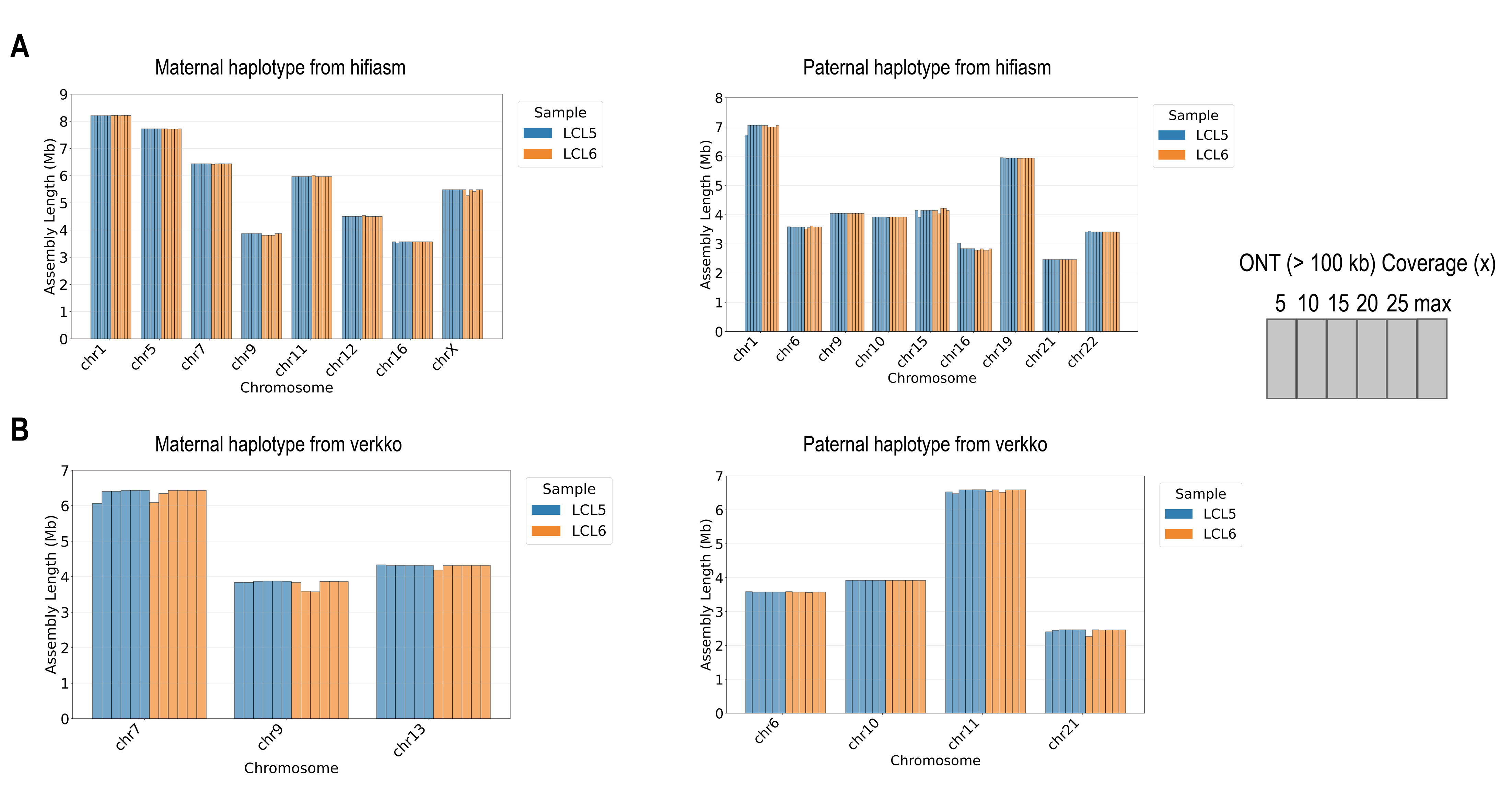
